# Ca^2+^ Plateau Potentials Reflect Cross-Theta Cortico-Hippocampal Input Dynamics and Acetylcholine for Rapid Formation of Efficient Place-Cell Code

**DOI:** 10.1101/2025.10.01.676159

**Authors:** Han-Ying Wang, Ying-Chieh Hsu, Hsuan-Pei Huang, Ching-Tsuey Chen, Xian-Bin Huang, Cheng-Ting Wang, Richard Naud, Ching-Lung Hsu

**Author notes:** Equal contribution (Ying-Chieh Hsu, Hsuan-Pei Huang, Ching-Tsuey Chen).

## Abstract

A central tenet of Systems Neuroscience lies in an understanding of memory and behavior through learning rules, but synaptic plasticity has rarely been shown to create functional single-neuron code in a causal and biophysically rooted manner. Behavioral Time-Scale Synaptic Plasticity (BTSP), identified *in vivo*, holds a great potential for explaining instantaneous hippocampal selectivity emergence by long-term potentiation (LTP), yet the cellular and endogenous mechanisms are unknown, impeding broader conceptualization of this novel rule for its algorithmic, systems-level and theoretical implications. Here, we addressed this gap by *in-vivo*, *ex-vivo*, *in-silico* and computational approaches to seek neurophysiologically inspired protocols for synaptically evoking Ca^2+^ plateau potentials and inducing potentiation in the CA1. We found induction of BTSP-LTP is best explained by a theta-oscillation-paced, gradually developed cellular state being supported with precisely timed weak ramping inputs. Remarkably, the previously presumed one-shot LTP for *in-vivo* place-field formation is possible under the influence of muscarinic activation. Through modeling, the notion of acetylcholine-gated BTSP gave rise to a computational advantage for low-interference continual learning. We further demonstrated that biophysics of Transient Receptor Potential (TRPM) and NMDA receptor (NMDAR) channels powerfully shapes the cross-theta dynamics underlying BTSP. These results which cover pre-, post-synaptic and neuromodulatory factors and their timing suggest fundamental principles for graded plateau potentials and hippocampal LTP induction. Overall, our work dissects cellular mechanisms potentially important for a prominent *in-vivo* hippocampal plasticity phenomenon, and offers a biological basis for framing BTSP as an input-dynamics-aware, neuromodulation-tuned synaptic algorithm.

## Introduction

Taking on central challenges for understanding brain dynamics requires explaining experience-dependent reorganization of neural circuits. Learning algorithms can be inferred from behaviors and psychophysics^1^. As a historic turning point, Donald Hebb’s seminal theory articulated how memories can be stored by changes of synaptic strength^2^, and the attention was shifted toward algorithms implemented by synaptic plasticity^3^. Nevertheless, despite extensive research on molecular/cellular mechanisms of hippocampal long-term potentiation (LTP)^4,5^, experimental analysis of its synaptic algorithms is rarely exercised in functional contexts like place-field formation, due to highly artificial induction parameters and protocol designs. On the other hand, elegant algorithms of functionally or theoretically constrained plasticity^6–11^ usually stay a distance away from biological implementations. In short, into half a century of synaptic plasticity work, explanations over adjacent biological and functional levels are still highly demanded to fulfill the promise of systems biology.

In this paper, our strategy for forging a function-implementation connection is to look into hippocampal plasticity *algorithms* from an empirical angle. To this end, we argue that a rational deconstruction of input stimulus patterns is urgently needed: such an algorithm shall be not only functionally useful, i.e., capable of producing robust formation of place cells, but also be informed by known physiological and biophysical constraints. We contend that, so far as physiological dynamics and states are relevant, biophysical specifics are more than “details” just for neural implementations; rather, they also shape and facilitate the sets of “rules” (or algorithms) crucial for LTP induction.

What is a good paradigm to pursue biophysics- and dynamics-informed algorithms, ideally under a rationale of principled stimulus design? Directly *in vivo*, experimenters “saw” the dynamics of membrane potential (Vm) immediately prior to the sudden formation of CA1 place fields, described as Behavioral Time-Scale Synaptic Plasticity (BTSP)^12,13^. The field-inducing event, a large, hundreds-milliseconds “Ca^2+^ plateau potential”, offers a unique perspective for algorithmic investigation into underlying presynaptic dynamics, postsynaptic messengers^14,15^ and neuromodulators given early dendritic integration research^16^. Unlike simple Hebbian Plasticity, widely seen as Spike-Timing-Dependent Plasticity (STDP)^17–19^ , which learns correlational structures of activation between cells via coincident presynaptic and postsynaptic activity, BTSP seemed to require no postsynaptic action potentials (APs) during presynaptic inputs. BTSP-LTP is thought to be recaptured in brain slices by evoking subthreshold EPSPs and a distinct dendritic Ca^2+^ plateau potential^20^ occurring within 1-2 seconds, under excitation of CA1 dendrites facilitated with intracellular Cs^+^. Except for cortical input^21^, however, the mechanisms are little understood, particularly regarding robust emergence of place cells triggered via the presumed synaptic potentiation.

Our synthetic experimental approach considering neuromodulation is in favor of the influential paradigms of instructive learning^20,22^. Acetylcholine signaling is important for cognitive and hippocampal mnemonic functions^23^; the fact that it modulates dendritic integration of cortical pyramidal cells^24^, plus some ideas of using neuromodulation for high-capacity learning in deep neural networks (DNNs)^25^, sets a strong prior seemingly natural to consider. In fact, any consolidation of a role for neuromodulators will fit with a broad family of rules under Three-Factor Learning^26^: activation of cortical synapses creates time-dependent traces of eligibility for their LTP induction, which is contingent on subsequent neuromodulatory input reporting behaviorally meaningful states. Together with the notion of circuit-state-dependent computation described above, these ideas constitute the organizing framework of hypothesis for our study.

Here, we built on cellular and biophysical mechanisms to construct a biologically plausible model for place-field-inducing, plateau-based LTP. As ion-channel biophysics is time- and state-dependent, the model instantiated an idea of internal states at CA3→CA1 synapses that continuously monitor input dynamics for inducing high learning-rate LTP. In line with the known role of acetylcholine for theta oscillations, theta and other dynamics of the inputs received by CA1 contributed to gradually developing strength of plateau potentials. Remarkably, the neuromodulator enabled one-shot induction of strong LTP, which supported learning across multiple environments while mitigating catastrophic forgetting in DNN models. Overall, synergistic combination of multiscale intracellular electrophysiology and experimental/theoretical approaches allowed us to rationally break down physiological parameters, which is otherwise impossible to dissect only in awake animals. We present a new view for effective brain-circuit learning algorithms.

## Results

### Rapid induction of hippocampal CA1 place-cell firing in goal-directed virtual navigation requires circuit inputs from external visual stimuli

We first asked that, in addition to excitatory inputs from the entorhinal cortex^21^, whether another major input from the CA3 could also play a role for the rapid induction of place-cell firing formation in the hippocampal CA1. Previous patch-clamp whole-cell recording for identifying place-cell formation in linear tracks for mice was carried out without active choice^12,21^. To engage synaptic plasticity possibly reflect information relevant to environments and goals, we implemented reward contingency in virtual-reality (VR), which required an active lick of head-fixed mice in a narrow reward zone (10 cm out of ∼170 cm) to trigger sucrose-solution delivery as the reward for each running lap (**Figure 1A**; see STAR Methods). This spatially accurate localization with potentially infinite behavioral options enabled efficient anticipatory lick patterns in the animals (**Figure 1B**). Consistent with previous work, a high proportion of tested CA1 pyramidal cells (∼79%; success = 11/14 neurons tested from 9 mice) developed a robust place field around the position where large Ca^2+^ plateau potentials were triggered, typically with no more than 2-3 times of brief somatic current injection (300-ms, +600-pA; **Figure 1C, D**).

**Figure 1.**
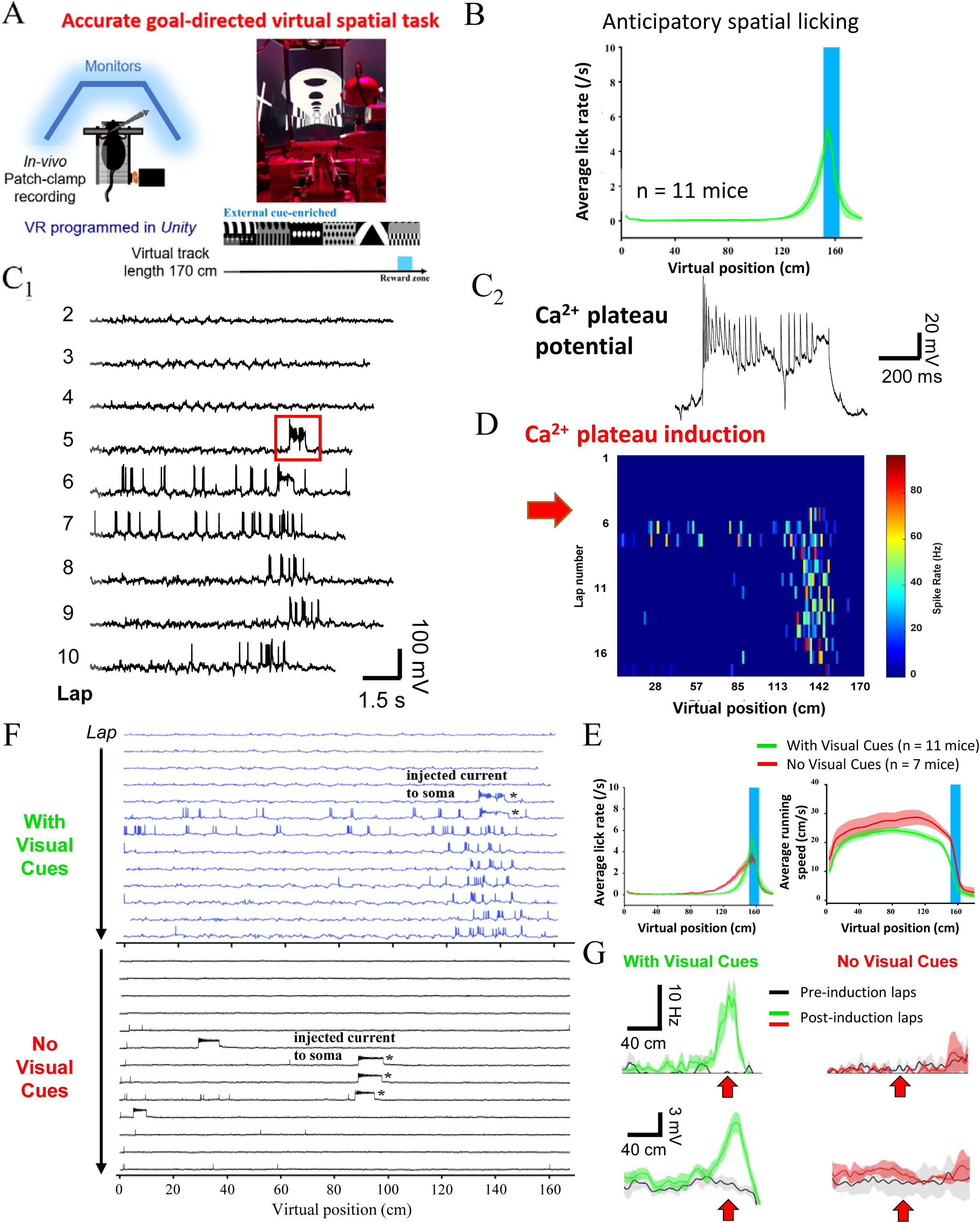
Rapid in-vivo formation of hippocampal CA1 place fields in goal-directed visual VR requires sensory stimuli. (A) Experimental setup of visually guided virtual reality (VR) for head-restrained mice, in combination with whole-cell patch-clamp recording from the hippocampal CA1. Delivery of rewards required active licks in the Reward Zone (RZ). (B) Mice licked in anticipation for the arrival of the RZ. (C) Brief somatic current injections of +300 pA and 600 ms in amplitude and duration triggered Ca^2+^ plateau potentials and rapid formation of CA1 place-field firing. (D) The heat map of lap-by-lap action-potential (AP) firing rate as a function of virtual position. (E) Both groups of mice under the normal VR condition and the VR condition without visual cues showed increase in lick rate and drop of running speed in anticipation of the RZ. (F) Representative example for lap-by-lap recording of Vm in mice under the normal VR condition or the VR condition without visual cues. The plateau induction protocol failed to evoke plateau potentials with regenerative spiking activity and place-field firing. (G) Induced suprathreshold firing and subthreshold Vm place field (i.e., synaptic ramp) and no apparent field induced in the VR with and without visual cues, respectively, from the mouse shown in **F**.

*In-vivo* and *ex-vivo* patch-clamp experiments suggested that CA3→CA1 inputs might play a role for BTSP computationally beyond just offering plain depolarizations. First, while the reduction of running speed and lick dynamics in anticipation for the reward both still existed (**Figure 1E**), a same level of somatic current injections failed to induce new place fields in the absence of visual cues (“No Visual Cues”) (0% success = 0/4 neurons tested from 3 mice; **Figure 1F, G**). Notably, in this condition, the amplitude of evoked somatic depolarizations was not different (With Visual Cues, 29.3 ± 1.8 mV, n = 14 cells; No Visual Cues, 35.2 ± 3.4 mV, n = 4 cells; p > 0.05), but they exhibited no apparent sign of regenerative Vm spiking (fast AP frequency during plateau induction, which is inversely correlated with regenerative Ca^2+^ spiking: With Visual Cues, 11.9 ± 2.3 Hz, n = 14 cells; No Visual Cues, 39.8 ± 10.3 Hz, n = 4 cells; p < 0.01, permutation test), different from what had been typically seen in the successfully induced cases with visual cues in the VR (“With Visual Cues”). Given our observation for clear relationships between successful LTP induction and Ca^2+^ spiking during plateaus in the experiments of acute brain slices (**Figure S1**), which highlighted an importance of regenerative activities for BTSP, these *in-vivo* results could be explained by a problem of generating effective forms of Ca^2+^/NMDA spikes. Because EC firing was not significantly affected in the lack of visual cues^27,28^, one possible scenario is that reduced firing rate of CA3 pyramidal cells (with their field locations unchanged) in visual-input deprivation^29^ led to a compromised condition for plateau generation and BTSP induction. This condition might more likely involve biophysical dynamics associated with the CA3→CA1 input, as we will unpack mechanistically in this paper, rather than simply a reduced total amount of proximal dendritic depolarizations.

### CA3 input dynamics shapes frequency-dependent BTSP potentiation kernels

Second, our data demonstrated that CA3 inputs contribute to induction of BTSP potentiation in a way sensitive to their firing dynamics. The effects of postsynaptic plateau potentials on BTSP are well established, but nearly no exploration has been conducted regarding presynaptic determinants. To parametrically test the presynaptic activity patterns in isolation from generation of plateaus, we used intracellular Cs^+^ in replacement of K^+^ to saturate plateaus (as in^13^) and varied presynaptic CA3 input firing controlled via extracellular field stimulation (**Figure 2A**). In our system, we induced very large LTP (**Figure 2B**) and reproduced the BTSP-LTP kernel with the backward and forward time constants identical to what had been initially reported (**Figure 2C**). BTSP kernels were remarkably sensitive to the firing frequency of CA3→CA1 (10 stiumuli, ranging from 10 to 100 Hz; **Figure 2D**). Interestingly, a range of frequencies (20-40 Hz) seemed to maximize the functional impacts of the LTP by having largest magnitudes of potentiation and time constants of the backward kernel (**Figure 2E**). Because the hippocampal CA3 fired at low rates *in vivo*^30^, this relatively low working range of CA3 firing level preferred by BTSP induction would suggest a potential functional optimization.

**Figure 2.**
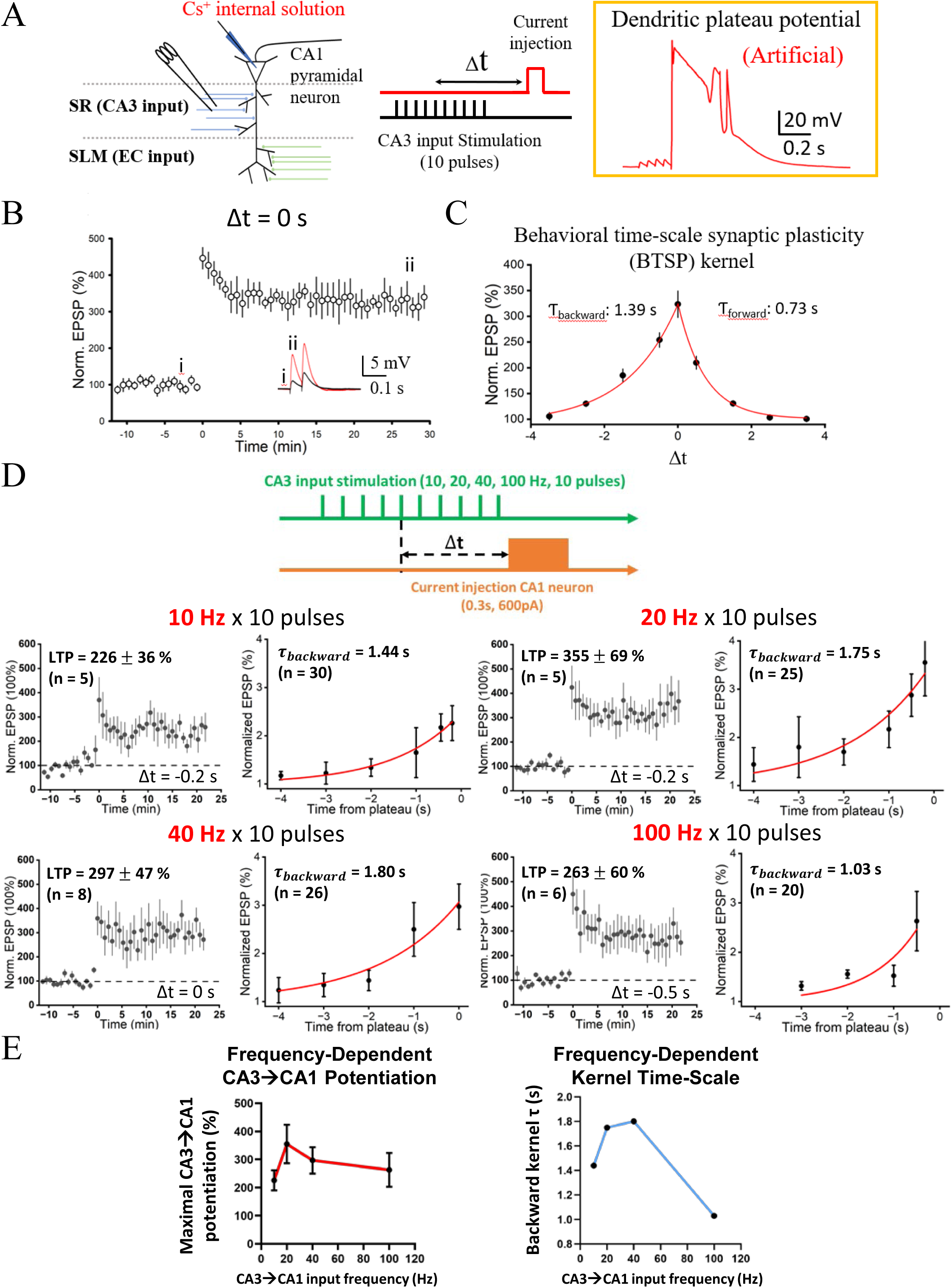
Smooth CA3-frequency-dependent BTSP potentiation kernel for CA3→CA1 synapses characterized with Cs^+^-enhanced saturated plateaus. (A) Experimental setup and a representative example of a plateau potential (per Bittner et al., 2017). (B) Normalized EPSP amplitude as a function of time with induction at Δt = 0 s. (C) The plasticity kernel, which was quantitatively nearly identical to Bittner et al. (2017). (D) Normalized EPSP amplitude as a function of time and backward kernels for LTP induced under different stimulation frequencies of the CA3 input. (E) Summary for maximal LTP magnitude for each CA3 input frequency and time constant for backward LTP kernel as a function of CA3 input frequency.

It is impossible to keep the duration of the entire stimulus pattern unchanged; in fact, the duration, frequency and number of stimuli are interdependent. To characterize the temporal features of relevant input-associated dynamics based on functionally unitary events, we applied a model-based method to derive the time constant of the underlying eligibility trace (ET)^14,31^ (**Figure S2A**). The Cs^+^ approach of the experiments (**Figure 2**) helped with model fitting toward a more restricted parameter space by achieving saturated plateaus, thus forcing an instructive signal (IS) as a fixed, unitary impulse (see STAR Methods). Another technical consideration was that presynaptic inputs and postsynaptic plateau activity could indeed interact at a level of their downstream second messengers. To approach the timescale of true temporally unitary ET events, we removed the data points with overlapped EPSPs and evoked plateau potentials from the kernels for subsequent fitting processes (**Figure S2B**). The outcome indicated that the ET time constant for presynaptic inputs was still largest around 20 Hz (**Figure S2C**). It showed that formal analyses based on biophysics-compliant signaling dynamics does not change the conclusion about the role of CA3→CA1 connections as informed by their temporal dynamics.

Besides the crucial role of EC input for BTSP indcution^21^, these observations argue for a place-field-forming rule that also take CA3 dynamics into account. Other than implying that CA3 inputs matter more, our results (here and below) highlight the potentially complex dendritic processing of synaptic dynamics for BTSP induction.

### Synaptically evoked large Ca^2+^ plateau potentials are sensitive to input dynamics of the circuits

Tackling the problem of BTSP cellular mechanisms hinged on an effective search for stimulus algorithms and activation patterns that trigger synaptically driven plateau potentials. To address this long-standing challenge, we sought physiological inspirations. First of all, decades of cellular electrophysiological studies have revealed that dendritic spiking is a thresholded phenomenon^32,33^. It is possible that plateau potentials require coactivation of many synapses to trigger^34^, so three stimulating field electrodes were placed in the *Statum Radiatum* (SR) and the *Stratum Lacunosum Moleculare* (SLM) to recruit three independent sets of synapses (**Figure S3**; see STAR Methods) across the dendritic domains of a CA1 pyramidal neuron (**Figure 3A**). We then considered physiologically relevant dynamic patterns to activate these synapses. One obvious target was theta rhythms^35^—it has been shown that theta-burst stimulation (TBS) induces dendritic Na^+^ spikes and Ca^2+^ plateaus and LTP^16,36^. Another good candidate was activation modulated by ramp frequencies^37^, which captures a prominent kinetic feature of place-dependent firing when the animal traversed spatial receptive fields. Ramp-modulated inputs can have strong influences on CA1 place-field responses^38^, input-output transformations, backpropagating action potentials, and dendritic Ca^2+^/NMDA spikes^39^.

**Figure 3.**
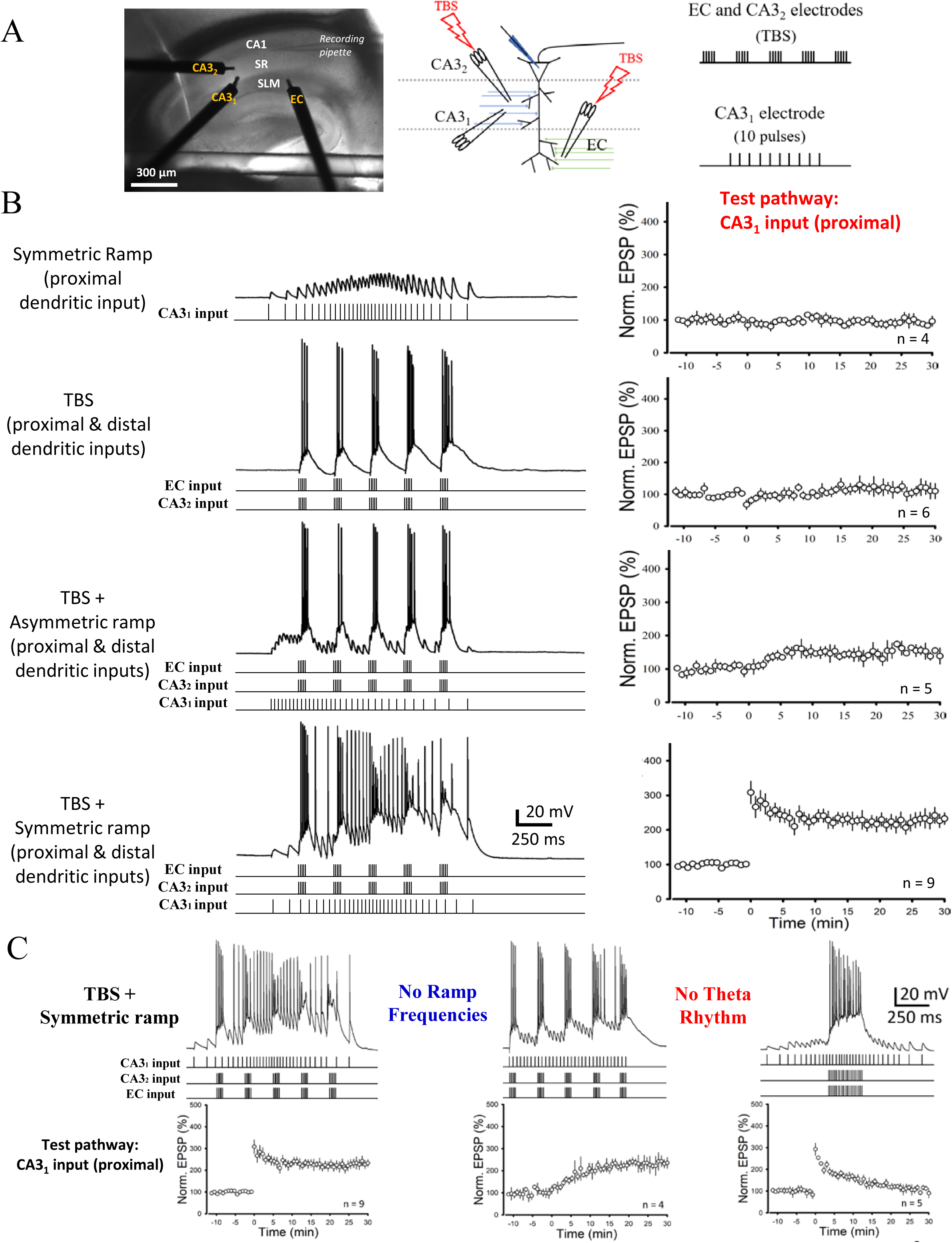
*Ex-vivo* rational design of stimulus patterns for synaptically evoked Ca^2+^ plateau potentials reveals a dependence on theta-rhythms and ramp dynamics. (A) Photo of an acute hippocampal slice with 3 field stimulating electrodes and 1 patch micropipette (*left*). Diagram for Experimental setup and the stimulus protocol. (B) Representative somatically recorded responses (*left*) and normalized EPSP amplitude as a function of time (*right*) in response to different stimulus protocols, progressively organized to have theta- and ramp-modulated dynamics from 3 independent sets of synaptic inputs. (C) Representative somatically recorded responses (*top*) and normalized EPSP amplitude as a function of time (*bottom*) in response to different stimulus protocols, with theta- or ramp-modulated dynamics removed from the protocols.

We rationally designed stimulus patterns and tested them hierarchically (**Figure 3B**). For each condition, the induction (or pairing) protocol was repeated only 5 times (see STAR Methods). Firstly, to mimic relatively weak CA3→CA1 synaptic activation in a silent place cell, which is believed to receive weak inputs tuned to random locations of the animal^39,40^, a ramp-modulated pattern (of 1.2-second duration and 50-Hz peak instantaneous frequency) was delivered to a test CA3 input pathway. The ramp activity alone did not produce LTP (**Figure 3B**, *first row*). When the ramp activity was combined with TBS delivered to a set of EC inputs, no LTP was induced at the test CA3 input either (n = 5 neurons; data not shown).

Takahashi and Magee (2009)^16^ showed that TBS simultaneously delivered to entorhinal cortical (EC) and CA3 inputs could triggered a form of dendritic Ca^2+^/NMDA potentials and LTP at the EC→CA1 synapses. Although it was not clear which dynamic properties specifically mediate the dendritic spiking in this case, this finding was an important prior. We therefore tested coactivation of EC and CA3 inputs by TBS; however, it failed to produce LTP at the test CA3 pathway (**Figure 3B**, *second row*). It is noteworthy that the test CA3 inputs would be located somehow in between the two TBS-activated pathways along the CA1 dendritic tree (**Figure 3A**), so this form of plateau potentials might be relatively weak, and not a global event^36^.

Finally, TBS of the EC and CA3 inputs was introduced with simultaneous activation of the independent test CA3 pathway in ramp dynamics. It is interesting to observe that a robust, immediate potentiation of test CA3 input was induced under this condition (**Figure 3B**, *fourth row*; LTP = 232 ± 20 %, n = 9 neurons), and it occurred with a symmetric ramp, rather than a “downward” (asymmetric) ramp (**Figure 3B**, *third row*), delivered to the test CA3 pathway.

There are some intriguing phenomenological parallels that can be compared against known *in-vivo* properties. Crucially, the theta- and ramp-modulated activities applied here was a rational “guess” for the patterns of what CA1 pyramidal cells “see” at their dendrites during active navigation, even for silent or non-place cells in which inputs of those patterns may be weak but exist^39^. EC and CA3 cells are engaged in theta oscillations during running and task engagements^41^, and some CA3 pyramidal cells have more place-firing to encode and transmit strong spatial representation^30^.

Remarkably, in response to co-occurrence of these circuit states, the resultant responses exhibited pronounced signatures of nonlinearities (**Figure S4A**)— reminiscent of somatic responses for large dendritic Ca^2+^ plateau potentials^12,16,42^— which were not observed in a condition with the same TBS state while a shorter 20-Hz train of stimulation was used for the test pathway, overlapping the 4^th^ and 5^th^ burst (when, in the TBS condition of EC and CA3 inputs, dendritically observable plateaus were known to initiate^16^). These characteristics include overall progressively increased spike half-width, decline of spike frequency, and appearance of shorter, broader spikes over the theta-cycles of the TBS; meanwhile, there was a slow Vm envelope growing over theta-cycles, most prominently after the 3^rd^ burst (**Figure S4B**). Notably, most of these effects had a clear, longer-lasting component between bursts of firing, a feature possibly involving after-depolarizing potentials (ADP) during plateaus^16^. The broader spikes may reflect dendritically initiated Ca^2+^ activities upon strong depolarization-induced inactivation of fast Na^+^ channels^43^, developing along with underlying, concurrent NMDA and persistent Na^+^ regenerativity^39,44^. Lastly, the immediate and strong potentiation of the CA3 input might underscore the plausibility for this form of plasticity to support hippocampal one-shot episodic encoding^17^.

### Properties of large synaptically evoked Ca^2+^ plateau potentials linked to algorithms and their biophysical basis

Having identified neurophysiologically inspired, rational design of a protocol permissive for triggering pronounced plateau potentials, we next took advantage of this well controlled system to construct educational speculation on the functionally relevant dynamic characteristics, which is essential to understanding the role of circuit states in efficient hippocampal learning. We altered the “TBS + symmetric ramp” stimulus pattern, which had been applied to 3 different synaptic pathways as inputs to the CA1. Here are the provoking results: theta-rhythms and ramp dynamics contributed to different aspects of the powerful, efficiently induced and early-(immediate) onset BTSP-LTP (**Figure 3C**). Removal of the theta rhythm (“No Theta Rhythm”) by collapsing the five bursts into one single train of 100-Hz stimulations completely abolished the sustained component of the LTP whereas a transient, early component stayed (*right*). In stark contrast, elimination of the ramp dynamics (“No Ramp Frequencies”) by applying stimulations to the test CA3 pathway at a constant frequency (∼38 Hz; same as the mean of the ramp used) disrupted the early, transient component of the potentiation while a slowly developing EPSP increase still constituted a sustained LTP (*middle*). In the latter part of the paper, our modeling work would suggest that significant supralinear summation of EPSPs remained in the No Ramp Frequencies condition, but nearly no supralinear summation was present in the No Theta Rhythm condition (see **Figure 8B** and **Figure S8**). We propose that supralinearity in the observed plateaus underlies the temporally persistent expression of BTSP-LTP.

Modeling induction processes offered a window for understanding BTSP algorithmically^14,31^. So far, Ca^2+^ plateaus have been postulated as an impulse function with invariant amplitude and duration, which then gives rise to downstream biochemical messengers in a form of low-pass-filtered signals, namely Instructive Signal (IS)^15^. Since the validity of this notion will lead to profound functional and computational consequences for BTSP, we would like to rigorously test the model assumption. With the data shown below, we are going to propose a broader conceptualization for plateau potentials, particularly under the physiological situations in which they can be driven partly by the same synaptic inputs to be potentiated (as we have demonstrated)—plateaus can reflect a continuously graded synaptic internal “state” for BTSP, more than (yet compatible with) a known role for offering discrete, “instruction”-bearing signals independent of ET.

We set out to directly test if a series of theta-entrained EC and CA3 inputs could bring a high state value for the “BTSP state”. That plateau signatures developing across theta-cycles (**Figure S4**) stimulated an intriguing speculation: the probability of BTSP induction for CA3→CA1 connections progressively increases upon entering the ramp and theta states. Experimentally, it was indeed the case since induced LTP became increasingly larger as the number of bursts in TBS, applied to one CA3 and the EC input, was changed from 1 to 5 (**Figure 4A**). The plateau-potential (slow underlying Vm envelope) area and other signatures emerged apparently corresponding to participation of the 4^th^ and 5^th^ burst, where induced potentiation also showed an abrupt enhancement, while the situation for only 3 bursts appeared to be intermediate (**Figure 4B**). This is consistent with the idea that dendritically initiated Ca^2+^/NMDA spiking tends to be more significant upon later cycles of continuous theta oscillations^16,40^.

**Figure 4.**
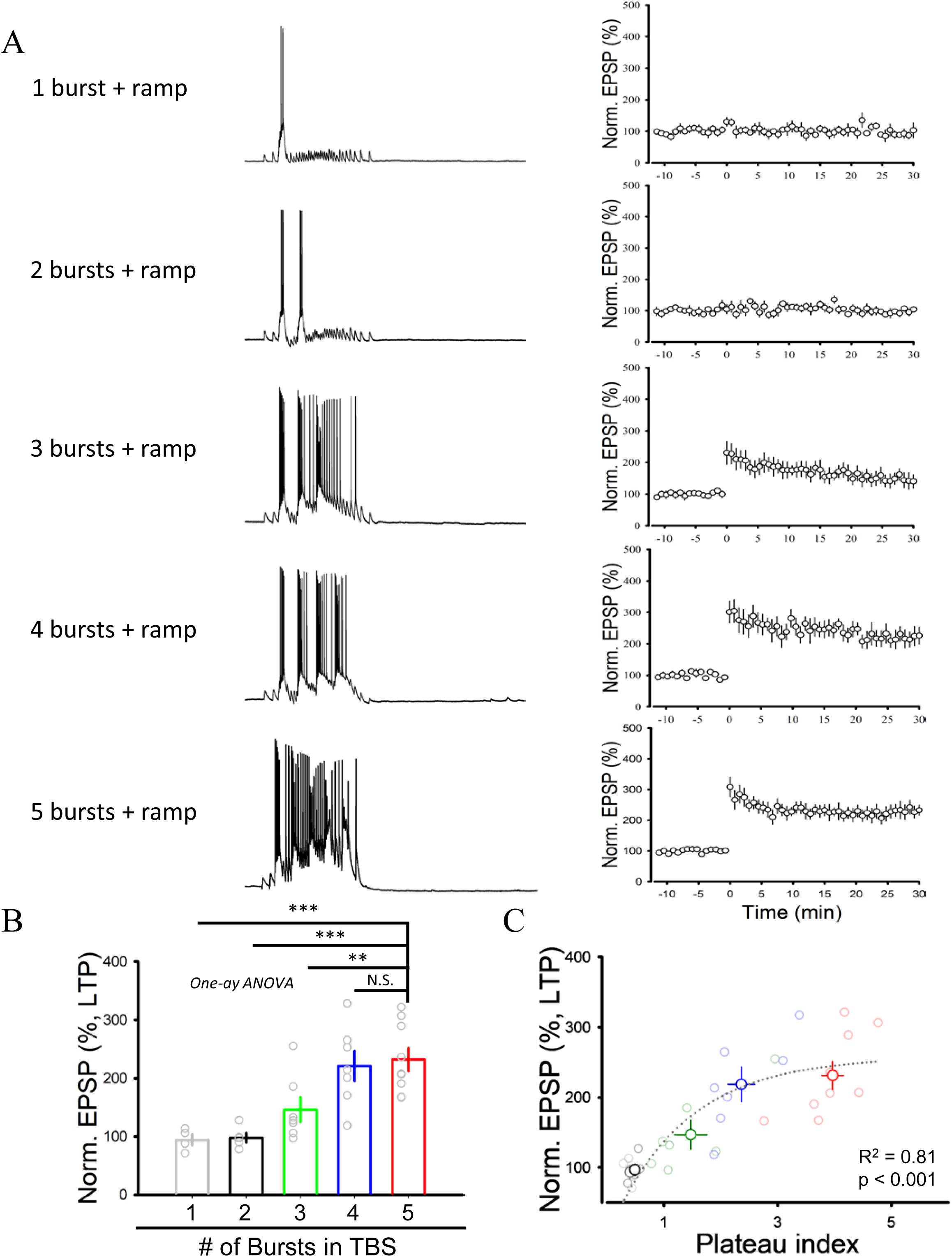
Time-dependent development of LTP-relevant plateau states driven by synaptic dynamics over theta cycles. (A) Representative somatically recorded responses (*left*) and normalized EPSP amplitude as a function of time (*right*) in response to a TBS pattern consisting of 1 to 5 burst stimuli. (B) Summary of the experiments in (A). LTP magnitude increased after TBS of 3 bursts, and most significantly upon TBS of 4 or 5 bursts. (C) Normalized EPSP amplitude as a function of normalized plateau area (EPSP integral from the 3^rd^ to the 5^th^ burst normalized with respect to 3-fold the integral of the 1^st^ burst), showing a sigmoid relationship.

Overall, these results demonstrated the properties of synaptically evoked large plateau potentials, which are consistent with a synaptic state variable in the CA1, sensitive to specific dynamic states of inputs from the upstream CA3 and EC, and representing LTP propensity over theta pacing. This paves a way for thinking of plateaus as a *BTSP state* capable of instantiating synaptic algorithms supported by their biophysics (see below).

### Plateaus as a synaptic state variable for BTSP induction is continuous

A conceptual usefulness of a state variable lies in the ability to encoding functionally relevant information in dynamic manners. The fact that plateaus were correlated with non-discrete LTP level (**Figures 4C, S5**) as well as able to compare inputs has suggested that plateaus operate as a continuous variable informing BTSP states.

To more directly test this idea, we envisioned a scenario wherein the synaptically evoked plateau condition could be continuously modified under exactly the same sets of inputs. Our implementation was to vary the time delay (Δt) between the test CA3 ramp input and TBS stimulation of EC/the other CA3 inputs, and we tried it first in a numerical model. Following the same formalism as Milstein-Romain^14^, and applying the ET time constants derived from our experimental measurements (**Figure S3**; additional experiments were performed for ramp CA3 input, ET time constant = 2.7 ± 0.8 sec; n = 25 cells), the test CA3 input generated ET, and the TBS of EC and CA3 input produced “IS”. In our experiments, TBS of CA3 and EC input (2 pathways only) could induce LTP at the CA3→CA1 synapses (LTP = 185 ± 22 %, n = 6 cells; *data not shown*), so we presumed coactivated CA3 and EC input triggered *weak* plateau potentials^16^. Note that it was still unclear whether this weak form of plateaus resulted in BTSP. Here, however, the “IS” resulting from such weak plateaus is named so only for convenience given that it does not reflect invariant plateaus anymore and can be modified via EPSP dynamics in the model. In other words, we use the term IS only for operational reasons, interpreting it as a hidden variable underlying *BTSP state* on top of its role for informing the *timing* of BTSP induction.

Through continuously modifying the condition of synaptically evoked plateaus via changing Δt, the computational model (**Figure S6**) supported that *BTSP state*, presumably reflected by plateau potentials, dictates LTP induction in a continuous domain. Three scenarios for IS (or *BTSP state* here) property was set up: Continuously Graded, Discrete and All-or-None. Inspired by the difference in plateaus observed between symmetric and asymmetric (downward) ramp condition (**Figure 3B**), the “IS” increased its strength continuously when Δt was within the time window of 0.5 seconds, and otherwise remained a constant as weak plateaus were considered effective for LTP induction here. In such Continuously Graded case, changing the time delay between test CA3 ramp input and TBS of EC+CA3 input produced a relatively smooth BTSP potentiation kernel (**Figure S6A**). On the other hand, in Discrete and

All-or-None case, the “IS” was modeled only as constants; however, weak plateaus were considered LTP-effective beyond 0.5 second in the Discrete case (thus the “IS” had two different non-zero values) yet not so in the All-or-None case (thus the “IS” was zero outside the 0.5-sec timescale). The latter two scenarios predicted BTSP kernels with different smoothness (**Figure S6B, S6C**).

Importantly, the actual experiments performed in acute slices (**Figure 5**) showed a smooth LTP kernel more consistent with the modeled Continuously Graded plateaus, which instantiated the synaptic state variable. Interestingly, LTP was still induced even at large Δt when no strong plateaus existed. The results critically demonstrate that synaptically evoked Ca^2+^ plateau potentials, in either a strong or a weak form, can dictate powerful synaptic potentiation with long time constants. Our experiments in reduced settings capture some defining characteristics of BTSP, with which the properties of synaptically driven kernels can be explained by systematically exploring the nature of underlying synaptic states.

**Figure 5.**
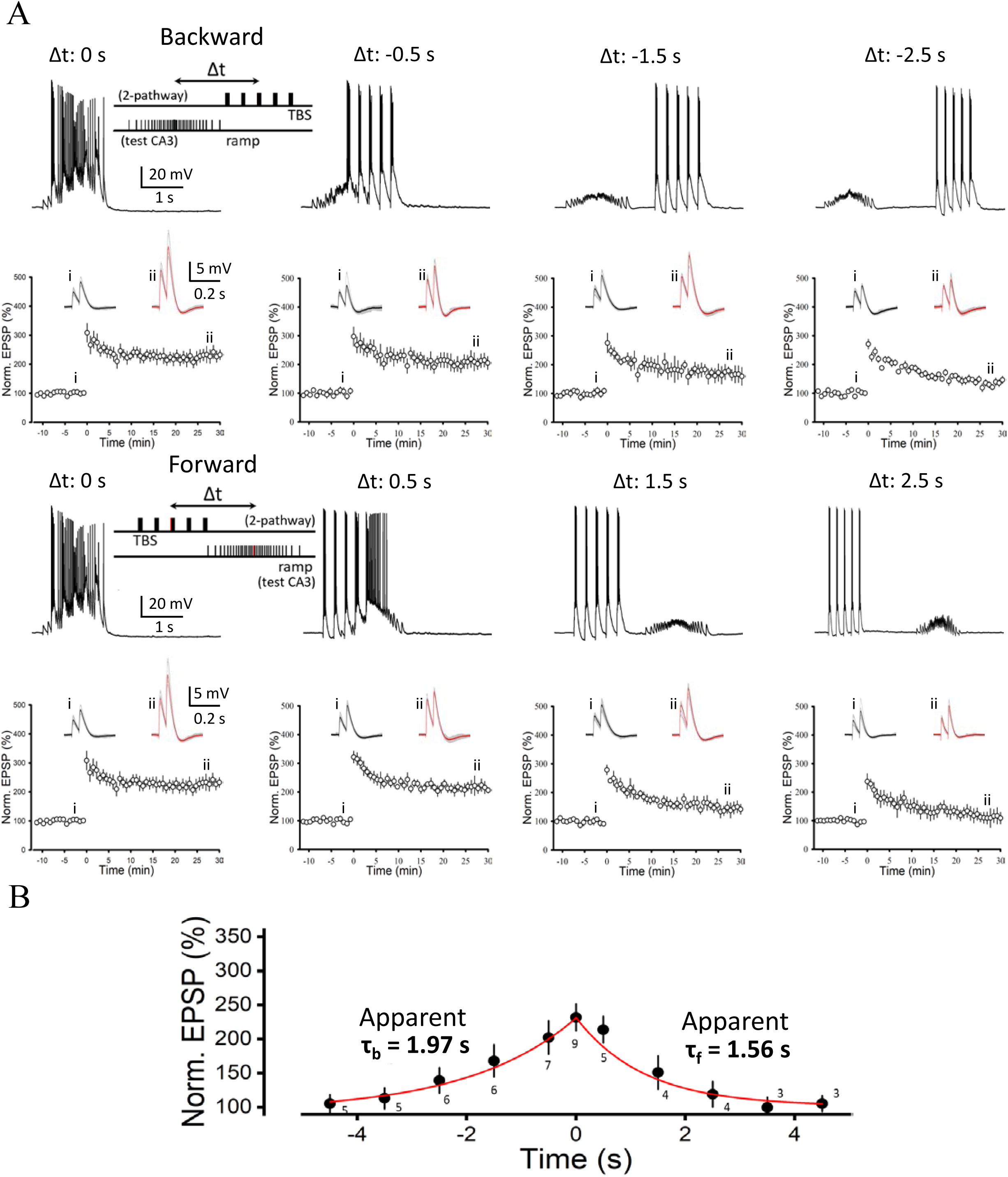
Varying plateau conditions through the timing of test CA3 ramp and theta entrained inputs generates smooth LTP kernel with BTSP signatures. (A) Representative somatically recorded responses (*top*) and normalized EPSP amplitude as a function of time (*bottom*) with example average EPSP traces before (i) and after (ii) LTP induction (*inset*) over different Δt. (B) LTP kernel summarizing the experiments shown in **A**. Note the comparably long time constants of the kernel (cf. **Figure S5**, *Continuously Graded Plateau*).

### Muscarinic cholinergic activation promotes one-shot BTSP and gates rapid formation of efficient spatial code

An important ingredient our theory lacks so far is a gating mechanism that controls BTSP. Induction of large LTP should be gatekept as, in the hippocampus, theta- and ramp-modulated dynamics are common at the circuit and synaptic levels^41,45^. Our work suggested that plateaus inform BTSP states over progression of theta oscillations (**Figures S4, 4**), and theta dynamics gates induction of the LTP (**Figure 3C**). Hypothesized to be gating signals for plasticity^24,26,46^, neuromodulators carry information about gross behavioral context. The medial septum, which contains acetylcholine (Ach)-releasing projection neurons, sends inputs to the CA1 and contributes to generation of theta-rhythms^23^. By critically controlling medial septal axons in the hippocampus, it was recently shown that theta power and sequences may underlie online spatial learning without involvement of replay^47^. Therefore, it was reasonable to hypothesize that one basis for theta-related gating is Ach signaling.

Endogenous release of acetylcholine could facilitate strong plateau potentials. Under the control of choline acetyltransferase (ChAT), channerhodopsin-YFP was expressed in medial-septum neurons through transgenics of a Cre mouse line (**Figure 6A_1_**).

**Figure 6.**
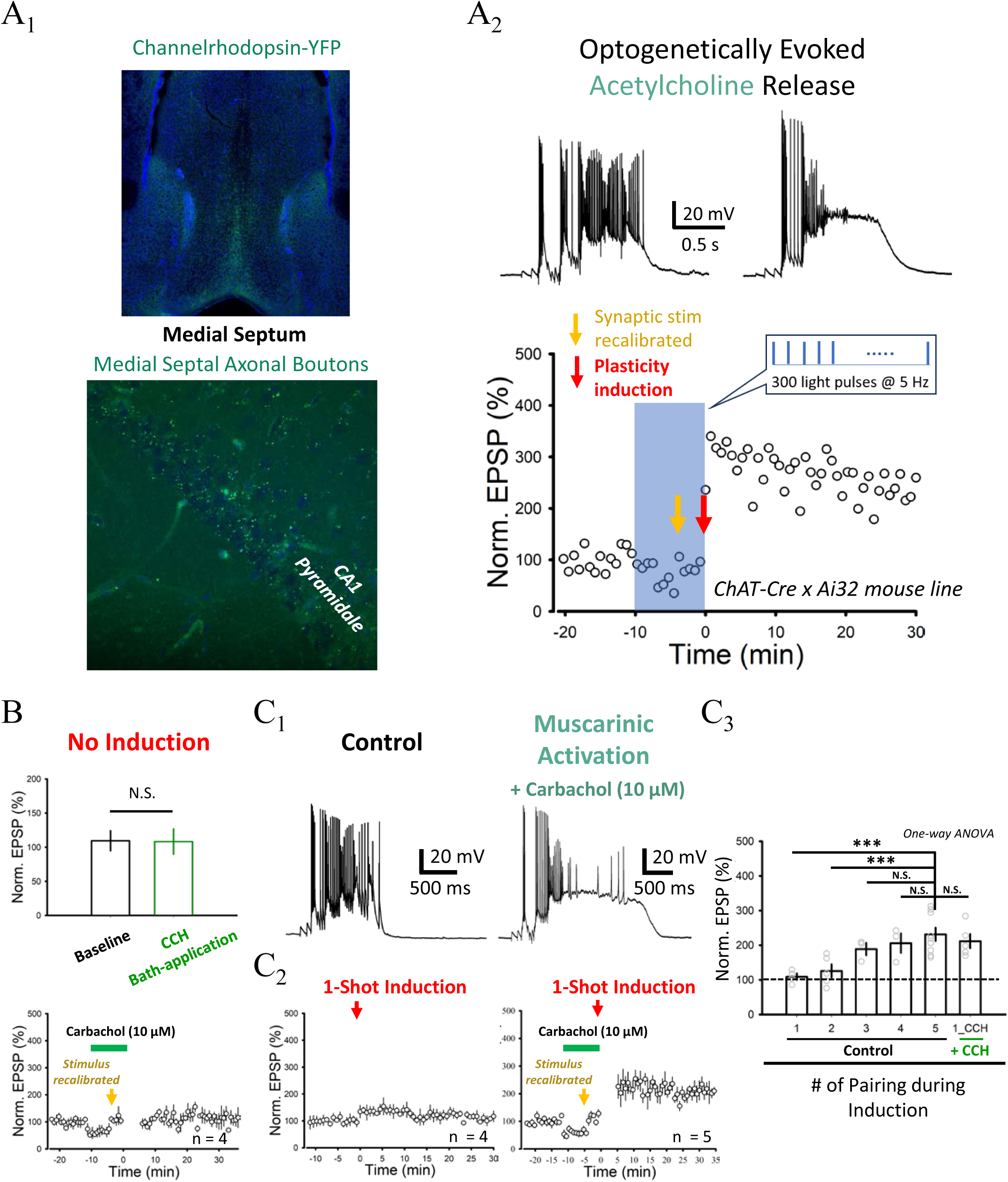
Acetylcholine release and muscarinic receptor activation promote plateaus and one-shot BTSP-LTP. (A) Representative confocal images showing expression of channelrhodopsin-YFP (green) in the medial septum and axons in and near the CA1 *Stratum Pyramidale* in ChAT-Cre x Ai32 mice (**A_1_**). Representative somatically recorded responses (*top*) and normalized EPSP amplitude as a function of time (*bottom*) with optogenetic stimulation of axons accessed under the active promoter of choline acetyltransferase (ChAT), with 300 brief light pulses at 5 Hz, during LTP induction (5 times; **A_2_**;). Note that the stimulus intensity for the test CA3 ramp input was adjusted during the period of optogenetic stimulation to compensate for the loss of dendritic excitation due to synaptic effects of acetylcholine signals (*brown/dark yellow arrow*; see STAR Methods). (B) *Bottom*, normalized EPSP amplitude as a function of time with activation of muscarinic acetylcholine receptors by carbachol (CCH) and no induction of LTP. This is an experiment controlling for longer-lasting effects of EPSP changes at the time 25-30 minutes after the washout, the time when LTP was evaluated for the experimental equivalent in **C**. *Top*, summary of the experiments indicating no change of EPSP amplitude once the stimulus intensity was adjusted (*brown/dark yellow arrow*). (C) Representative somatically recorded responses (**C_1_**) and normalized EPSP amplitude as a function of time (**C_2_**) for 3-pathway LTP induction (1 time only; i.e., “1-shot induction”) without or with activation of muscarinic acetylcholine receptors by carbachol (CCH). Summary of the experiments indicating that muscarinic activation increased the rate of synaptic weight update (or lowered the threshold for LTP), allowing BTSP-LTP of the same magnitude to be induced by 1-shot induction alone (**C_3_**).

Optogenetic excitation of medial septal axons was delivered at a theta-rhythm (5 Hz). While it was clear that synaptically evoked plateaus could be enhanced in response to the opto-stimulation (normalized plateau integral: 4.79 ± 0.63, n = 2 cells; **Figure 6A_2_**), induced LTP was not significantly larger under the current sample size (LTP = 230 ± 11%, n = 2 neurons). Nevertheless, the antagonist for muscarinic Ach receptor Atropine (1 μM) significantly reduced the plateau area (normalized plateau integral: Control, 3.85 ± 0.23, n = 9 cells; Atropine, 2.99 ± 0.32, n = 7 cells) and LTP (Atropine, LTP = 164 ± 17%, n = 7 cells; p < 0.05, 1-way ANOVA with *post-hoc* means comparison), indicating a role of endogenous, synaptically release acetylcholine for BTSP.

Probing the other direction of gating by further activation of muscarinic receptors, beyond the synaptically release regime in acute hippocampal slices, can be achieved by exogenous application of agonists. We bath-applied Carbachol^24,48^ during the induction (10 μM; for no more than 10-12 minutes to avoid irreversible effects^49^, see STAR Methods for details) when the stimulus intensities were readjusted (according to somatically measured EPSP amplitudes) to compensate for the temporary loss of dendritic excitations due to reduction of EPSPs (see STAR Methods). Importantly, a matched design of the experiment was performed, while the LTP induction was omitted, to confirm that no lasting effects of EPSP changes occurred at the time when we evaluated LTP level (**Figure 6B**) under this pharmacological condition. In response to the pairing protocol, we observed significantly enhanced and prolonged plateau potentials (**Figure 6C_1_**; normalized plateau integral: Control, 3.85 ± 0.23, n = 9 cells; Carbachol, 6.52 ± 0.52, n = 5 cells; p < 0.001, Student’s t-test); in this case, only 1 pairing was sufficient to trigger LTP of the same magnitude (**Figure 6C_2_**; LTP = 212 ± 21%, n = 5 cells; p > 0.05, 1-way ANOVA with *post-hoc* means comparison). In the control (no CCH) condition, multiple pairings were needed to generate large and rapidly induced LTP, a requirement was eliminated by activation of muscarinic Ach receptors (**Figure 6C_3_**). As the consequence, acetylcholine signaling may convert the few-shot plasticity induced by synaptically evoked plateau potentials into a one-shot plasticity. This observation makes hippocampal one-shot plasticity a result of a tripartite conjunction of distal dendritic input, perisomatic excitation and cholinergic signaling.

We also examined the effects of pharmacological inhibition of the medial septum (MS) on place-field formation with the experiment of BTSP induction *in vivo* (see **Figure 1**) when 3.2 mM Muscimol (GABA_A_ receptor channel agonist) was stereotaxically injected prior to the session of whole-cell recording in VR. Currently, unlike the control condition in which place fields were rapidly induced in the contralateral hippocampal CA1 under saline injected to the MS (n = 2 cells), the fraction of theta power was lowered and the CA1 cell failed to express new place fields in response to induction with Muscimol injected to the MS (*data not shown*). As this is only preliminary, more work will be conducted to confirm the observation.

We contemplate that this is a conceptually important finding, as neuromodulators have been long postulated as a critical scalar feedback signal which gates LTP for appropriate synaptic credit assignment in deep neural networks^22,46,50–52^. However, direct evidence for effective modification of synaptic update efficiency is scarce^26,53^. Our findings lead to a theoretical possibility that regulated and sparse LTP events with high-learning rates can be enabled by such scalar feedback and the modulation of BTSP state variables. We explored this idea in a computational model.

To illustrate how control of the strength of plasticity by neuromodulation can be beneficial, we simulated place-field formation and its maintenance in CA1 neurons through multiple environments. Following the model by Bittner et al.^13^, we modeled CA3 neurons as having place fields before those to emerge in the CA1 after plateau induction. Plateau induction was further modeled as arising from a place-field-like teaching signal originating, at least, in part from the EC. We extended this model in two main aspects: (1) To capture place-field formation for different environments: it was done by providing different place fields to CA3 neurons, and presenting different environments in stages (**Figure 7A**). (2) To capture the effect of neuromodulators on plasticity: focusing on ACh (or norepinephrine, NE), which can signal aspects of novelty, we scaled the learning rate by a factor that was high when the environment was seen for the first time and low when it had been seen many times (**Figure 7B, C**).

**Figure 7.**
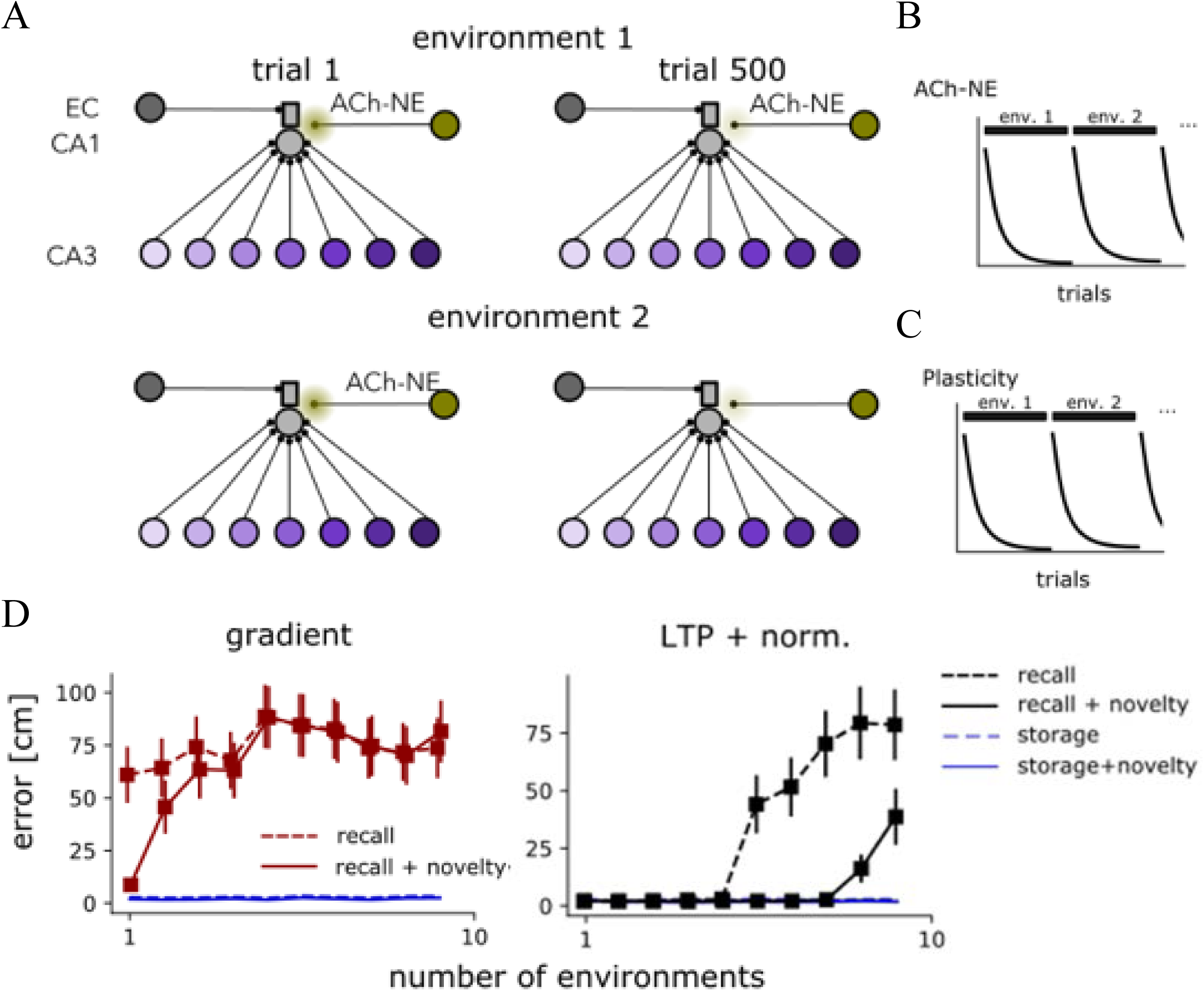
Neuromodulator-facilitated one-shot LTP can enhance memory capacity upon continual learning over many environments. Modeling the neuromodulation of plateau duration and synaptic plasticity for memory storage and recall. (A) Schematic of the modeled network showing entorhinal cortex (EC) input communicating a target signal. CA3 units represented different positions in space (color coded). A neuromodulation was provided by acetylcholine or norepinephrine (ACh-NE), which was potent on the first trial and decreased with experiences over trials, representing a novelty signal. New environments were modeled by scrambling the association between cells and space (*bottom row*). (B) A graph showing the modeled scheduling of ACh-NE release as the agent was exposed repeatedly to the same environment and the switches to a new environment. (C) As in **B**, but for the net amount of plasticity induced by plateaus. (D) Storage and recall error as a function of the number of environments seen by the agent. The presence of novelty signal (ACh-NE) made a significant difference in recall error (but not in storage error, so the training process remained the same), showing that the recall was improved with memory interference limited by sparse, one-shot-like BTSP-LTP learning events gated by the neuromodulation (*right*); the effect was markedly reduced in the case of gradient descent method for updating the network.

**Figure 8.**
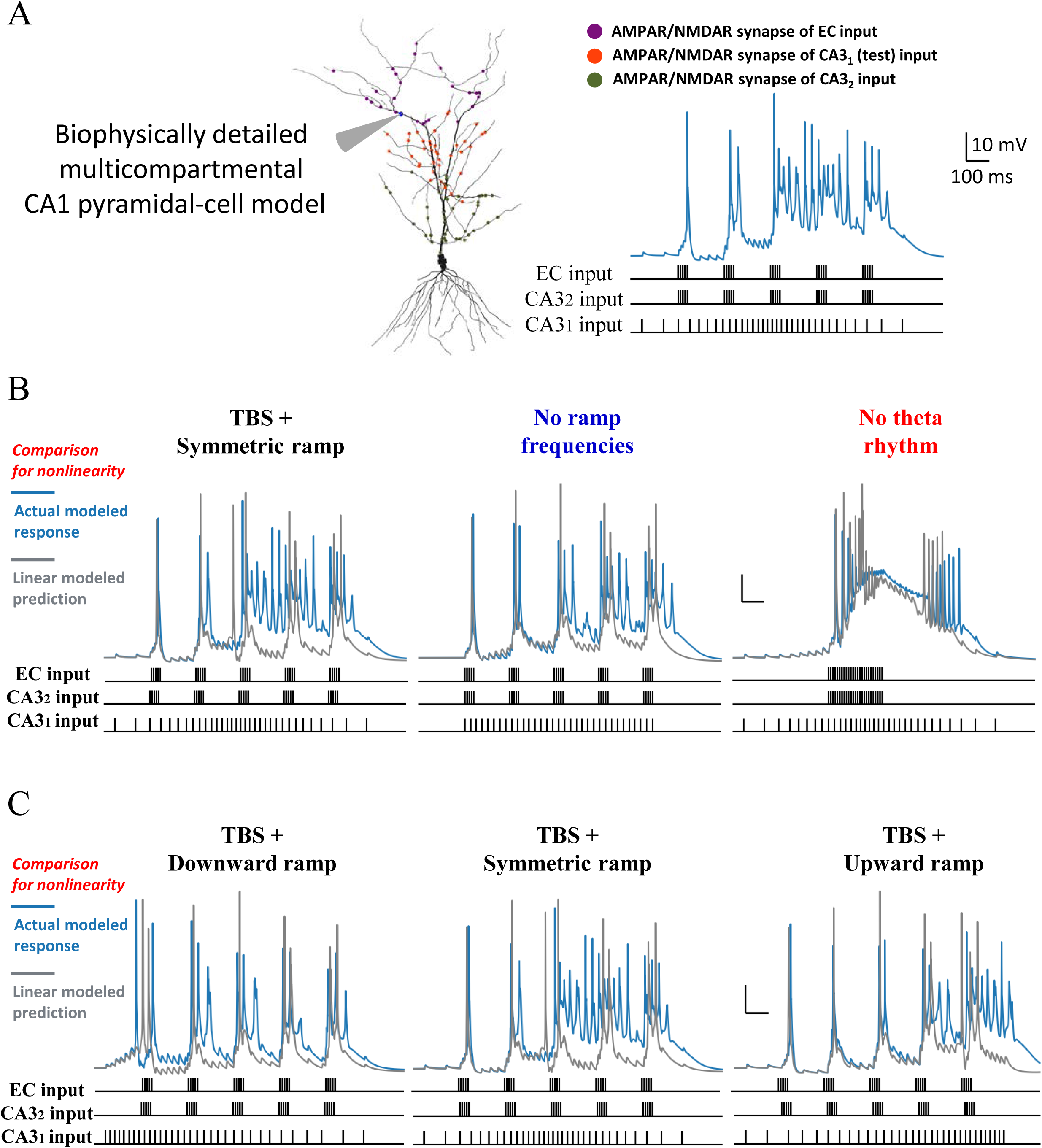
Emergence of synaptic plateau dependence on ramp and theta-cycle timing from realistic ion-channel biophysics of CA1 pyramidal neurons. (A) *Left*, model setup. A biophysically realistic multicompartmental model of hippocampal CA1 pyramidal neurons, containing 12 different ion-channel conductances and constrained by multiple independent experimental data, including our own experiments. Morphology and the spatial distribution of synapses along the dendrites were based on array-tomography data from Bloss et al. (2018). *Right*, 3 distributed, independent sets of AMPA/NMDA synapses were activated with the theta-burst- and ramp-dynamics, resulting in plateau potentials gradually developing over theta cycles, with increased spike width and moderately reduced spike frequency over time. (B) The model predicted (without specific tuning) the responses to *No ramp frequencies* and *No theta rhythm* condition well. Note that these responses were not part of the parameter optimization process during mode fitting. Linear sum of the responses to the individual stimulus applied for each input pathway (gray line) was compared with the observed response to the full stimuli applied to all pathways (blue) in order to assess the supralinearity of the outcome in the model. The theta pacing appeared to contribute to the surpralinear response much more than the particular dynamics of the ramp pattern. (C) The model also predicted that the plateau response was stronger in *TBS + Symmetric ramp* than *TBS + Downward ramp* (Figure 3). The even more pronounced supralinear plateau in the *TBS + Upward ramp* condition suggests that, for an ongoing, cross-theta state transition made into dendritic plateauing, it is the timing for the high-firing part of the ramp, instead of ramp symmetry, which is more important for plateau quality and accelerating the response supralinearity.

This model captured an enhanced LTP in novel environments. Computing the storage and recall error for decoding space as a function of the number of different environments, we found that adding the novelty signal supported by our experimental observations increased the number of environments that can be effectively decoded, before leading to a significant increase in the recall error (**Figure 7D**; *right*). As opposed to the similar situation but done with traditional gradient descent (*left*), this effect is to be expected since the hypothesized novelty signal effectively constrains the LTP to the moments where it is needed for relevant new place-field formation, reducing the total amount of less specific plasticity undergone in the network. Since the storage error (computed during training) had remained low, we conclude that a novelty signal that tunes BTSP-LTP reduces forgetting without affecting learning.

In conclusion, our results may open a new avenue for consolidating a role of neuromodulation in the algorithms for synaptically evoked plateau potentials and highly efficient plasticity, by revealing a mechanistic link between acetylcholine signaling and the proposed BTSP states. One-shot BTSP might enable robust place-cell code for hippocampal episodic memories.

### A biophysically realistic dendritic Ca^2+^ plateau model predicts the dynamic properties of plateaus observed in experiments

Biological implementations may shape the properties of neural algorithms. Finally, we performed additional pharmacological experiments, and built a realist model of CA1 pyramidal neurons to understand the biophysical mechanisms.

The CA1 model was adapted from the existing predecessors based on Cembrowski^44^ and Hsu^28^. Fundamentally, none of the already existing multicompartmental models would be able to recapitulate hippocampal Ca^2+^ plateau potentials (even as one defined in Takahashi and Magee^16^), so our critical contribution of creating realistic models for synaptically evoked plateau potentials is highlighted below (see STAR Methods for details; **Figure S7**): (a) a good diversity of voltage-dependent Ca^2+^ channels were incorporated and constrained according to the data of dendritic Ca^2+^ spikes, which the models of dendritic integration had often fell short of; (b) calcium-dependent potassium (BK and SK) channels were appropriately used to overcome the problems regarding over-excitations and uncontrolled AP waveforms, typically encountered during modeling of strong forms of dendritic Ca^2+^/NMDA potentials (*personal communication*, Aaron Milstein); (c) while most practices constrained the model only in terms of branch-level integration or dendritic Ca^2+^ spikes in response to current injected to the apical trunk, we tuned several forms of Na^+^- and Ca^2+^/NMDA-based nonlinearities simultaneously, which led to a balanced recreation of CA1 dendritic integration, cable properties and the capability of 3-pathway dendritic plateau potentials (**Figure 8A**); (d) we introduced Melastatin-like Transient Receptor Potential (TRPM) channels^38^, which accounted for the slow Ca^2+^-dependent build-up of plateaus and thus the state value for BTSP induction (see below).

Our model generated predictions regarding sensitivity to the input dynamics of the EC and CA3, remarkably in agreement with the experimental results (see **Figure 3**). First of all, the simulated responses to *No ramp frequencies* had a reduced plateau area while the response to *No theta rhythm* showed a single bout of depolarization. When these responses were compared against the linear sum of the simulated responses to each individual input (EC or one CA3 only), it was clear that *TBS + Symmetric ramp* condition exhibited the most supralinearity, followed by *No ramp frequencies* and *No theta rhythm*, in this order (**Figures 8B, S8**). As mentioned earlier, with **Figure 3C** considered together, this result supported that supralinear responses of apparent plateau potentials contribute to the induction of the sustained component of BTSP-LTP. Interestingly, without any extra tuning, the model precisely predicted a much reduced supralinearity for the condition of asymmetric (downward) ramp (**Figure 8C**; see **Figure 3B**). This prompted us to further consider the role of ramp dynamics mechanistically, by changing the asymmetry of the CA3 ramp and shifting the centroid of activity to later times in the model (**Figure 8C**, *right*). The even more pronounced plateau supralinearity observed for the later times suggested that it is the relative timing of the high-frequency firing of a ramp with respect to an ongoing theta rhythmicity which determines the level of plateau generation. This timing-dependent effects echo a central theme of our argument in relation to synaptic internal state: the theta oscillation of upstream inputs should be viewed to be associated with a dynamic state variable, in this case again relevant to BTSP; these EC/CA3 states can be read out as part of the *BTSP state* reflected by plateaus, then instructing robust LTP.

The realistic CA1 model offered a rich array of simulated outcomes that assisted with the interpretations about the underlying ion-channel mechanisms. Most significantly, the dependence of plateaus initiated in the later part of theta oscillation on NMDA-type glutamate receptor (NMDAR) channels, voltage-gated Na^+^ channels (VGSCs, most likely persistent Na^+^ current^39^), TRPM channels and R-type and L-type voltage-gated Ca^2+^ channels (VGCCs), as well as short-term synaptic release probability (P_r_) provides insight on relevant biophysics (**Figures 9A, S9**). The dependence on NMDARs and VGSCs naturally emerged in the model, with the former in agreement with the pharmacological results obtained with 50 μM APV for blocking NMDARs (**Figure 9B**). In this set of experiments, notably, the synaptically evoked responses in *TBS + Symmetric ramp* and *No ramp frequencies* condition showed more apparent sensitivity to the blockade of APV, which was correlated with the magnitude of the inferred sustained (persistent) component of BTSP-LTP across conditions. In contrast, the apparent APV sensitivity had no correlation with the early, immediate potentiation of the BTSP. It corroborated our interpretation that the theta-dependent plateau supralinearity, mostly mediated by slower NMDAR channels (**Figure S9**), is the major contributor (likely via Ca^2+^ influx) to the long-lasting potentiation of BTSP.

**Figure 9.**
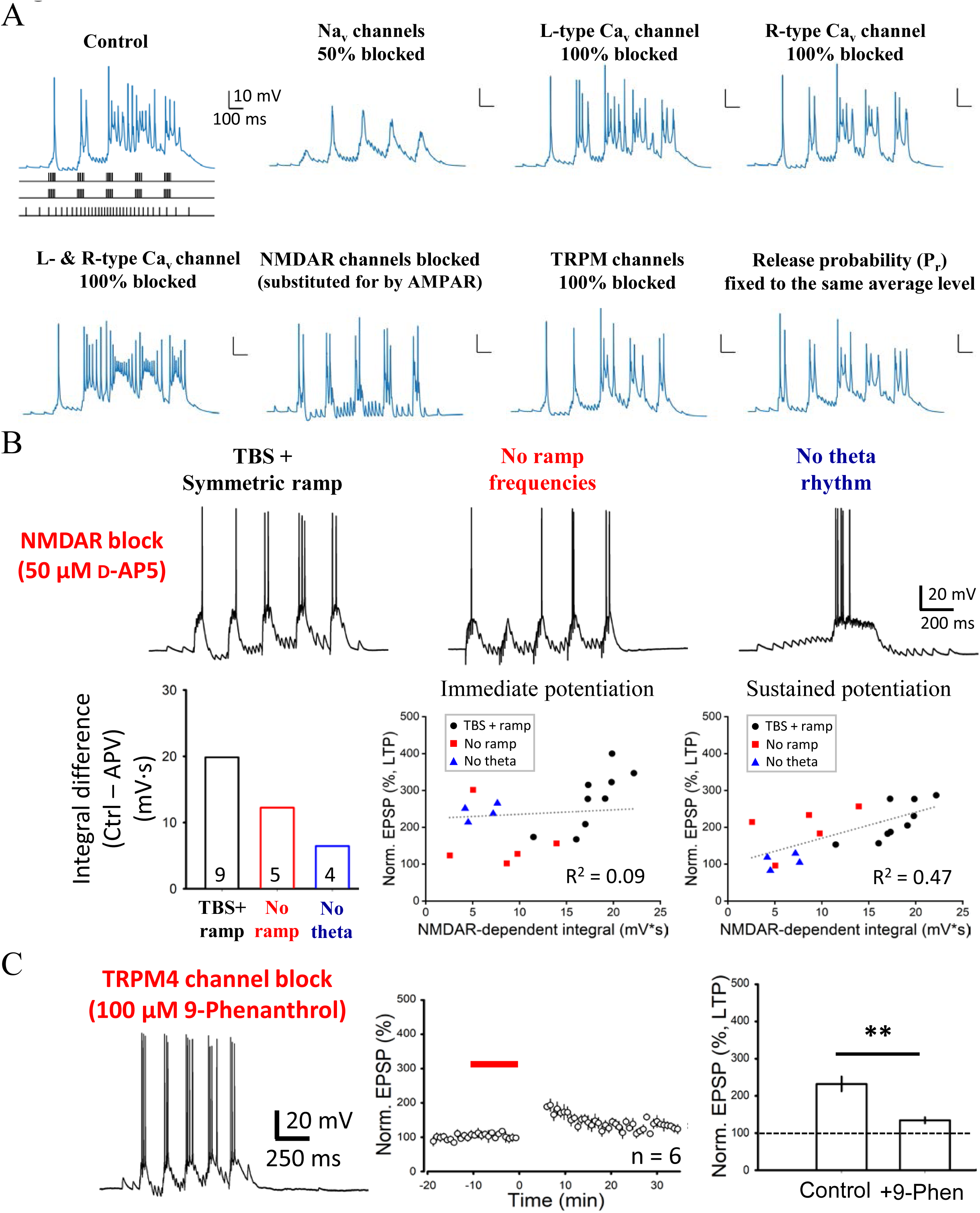
TRPM, NMDAR channels and short-term synaptic properties underlie time-dependent plateau generation across theta cycles. (A) (Modeling) Dendritically recorded response to EC and CA3 inputs with theta- and ramp-dynamics in control, simulated pharmacology and manipulation of presynaptic release probability (Pr). Note that broad Ca^2+^ spikes emerged in response to 50% block of voltage-gated sodium channels (for impairing persistent Na^+^ current and initiation of dendritic Na^+^ spikes without affecting synaptic transmission; Kim*, Hsu* et al., 2015; Hsu et al., 2018), which is consistent with actual experimental observations; therefore, the effect of persistent Na^+^ current on the slow, cross-theta EPSP depolarizations was more than it appeared. (B) (Experiments) *Top*, representative somatically recorded response to EC and CA3 inputs with theta- and ramp-dynamics in control (*TBS + Symmetric ramp*), *No ramp frequencies* and *No theta rhythm* condition, with the NMDAR channel antagonist 50 μM D-AP5 applied in the bath during induction. *Bottom*, using parallel sets of between-group control as a reference, the difference in EPSP integral was calculated, of which the LTP level (Immediate potentiation: 0-3 min post-induction; Sustained potentiation: 25.5-30 min post-induction) of each condition was plotted as a function. Compared to the early, immediate BTSP potentiation, the result suggested that the late, sustained BTSP potentiation of the ramp CA3 input is more sensitive to the activation of NMDAR channels. (C) (Experiments) Similar to B but with the TRPM4 channel blocker 100 μM 9-Phenanthrol applied in the bath during induction. Summary of experiments showing that BTSP-LTP was nearly abolished.

The observed plateau dep endence on TRPM channels, R-type VGCCs and Pr was not spontaneous predictions made from the original model. Specific experiments were performed for relevant measurements regarding TRPM (**Figures 9C**, **S10A-C**), R-type VGCCs (**Figure S10D**), L-type VGCCs (**Figure S10E**) and Pr (^36^, see STAR Methods for details). Regardless of an emerging property or a fitting target, simulated dependence on these mechanisms offered mechanistic plausibility for slow activation (NMDAR channels and R-type VGCCs), nearly non-inactivating kinetics (persistent Na^+^ current) and cross-theta facilitation of synaptic release (Pr) to contribute to plateaus. Note that some broad Ca^2+^ spikes appeared in simulated 50% Na^+^-conductance block, a previously known effect^55^ that was independent of overall reduction of the late-phase plateaus. It is also noteworthy that R-type and L-type VGCCs had a strong contribution to BTSP induction (**Figure S10D, E**) despite a relatively weak effect on plateaus compared to NMDARs (**Figure 9B**) and TRPM4 (**Figures S10A, B**; cf. **Figures 9A**, **S9** for the model), suggesting a possibility that R-type VGCCs provide biochemical second messengers (Ca^2+^) other than simply electrogenesis for plateaus.

TRPM-channel opening kinetics are voltage- and calcium-dependent^38^; to our surprise, this slow conductance only recently appreciated in the CA1 made an incredibly significant contribution to plateau area (cf. **Figure S9B** for the model) and BTSP potentiation, particularly the TRPM4 subtype, rather than TRPC channels (**Figure S10A-C**). These remarkable effects could be explained by the postsynaptic nanodomain Ca^2+^ dynamics with a slow decay (hundreds of milliseconds)^38^ and the Ca^2+^-dependent, non-inactivating kinetics of the TRPM (**Figure S11**). This ‘molecular reader’ for synaptic states is an intriguing discovery that potentially helps explain both the mechanisms and algorithms underlying plateaus and the associated *BTSP state*. Altogether, we think that synapse-state-dependent plateau generation is an elegant consequence of a sequence of biophysical events: arising from concerted actions between moderately slow ion conductances that sense excitatory synaptic inputs, sustained channel availability via theta ‘pausing’, electrophysiological state matching by temporally biased ramp dynamics, and a cross-theta molecular readout of TRPM linking to the overall state of BTSP induction. Furthermore, given the known cholinergic modulation of TRPM4 channels^56^, this biophysical implementation could connect to the stated scalar feedback and gating mediated by acetylcholine signaling (**Figures 6**, **7**) and elucidate a basis for coupling neuromodulation to BTSP state variables.

## Discussion

Decades of research on synaptic plasticity have found LTP of many forms and mechanisms^4,20^, but, as the majority of the knowledge came *in vitro* (*ex vivo*), a core challenge is how they causally as well as algorithmically relate to experience-dependent neural dynamics and behaviors. The hippocampus is a central model for synaptic plasticity for various reasons, especially its recognized roles in rapid and efficient code for episodic memories^57,58^. Our project attempted to address this fundamental problem by experimentally determining the biophysically and neurophysiologically principled mechanisms of BTSP, and, assisted with computational techniques, bridging this understanding with theoretically relevant plasticity algorithms, hopefully unraveling its aspects of behavioral functions in the future.

BTSP is at the center of our interest because it is the only putative LTP form we know which unambiguously generates single-cell selectivity for receptive fields in the awake, functioning brain^20^. On basis of whole-cell patch-clamp recording performed in the CA1, the theory for BTSP mechanisms had strong and rational priors—an induction through conjunctive activation of excitatory inputs from the CA3 and ECIII^12^. Nevertheless, a huge gap has existed between the efficient formation of single place-cell code and the specific mechanistic details (and thus the actual algorithms this plasticity rule implements), as most BTSP properties were indirectly inferred *in vivo*^14^. With patch clamp in highly controlled systems as well as rational design of synaptic stimulus patterns, the major threads of our findings are: (1) we envisioned that EC and CA3 inputs were both important to set the biophysical “thresholds” for LTP-inducing, regenerative dendritic spiking, and used two potentially highly prominent place-firing-related dynamics in the hippocampus, theta-rhythms and frequency ramps, to construct induction patterns; (2) we found the recipe of theta- and ramp-modulated inputs could trigger plateau potentials accompanied by robust and immediate expression of large LTP, which had signatures of BTSP; (3) we leveraged what experiments in brain slices can best offer, analyzing the properties of BTSP induction associated with the dynamics of inputs; (4) the results highlighting the importance of theta and the input timing in a train of theta oscillations, together with the smooth kernels of BTSP-LTP as the result of involving both strong and weak forms of plateaus, led us to postulate plateaus as an electrophysiological manifestation of continuous “hidden” state variables for BTSP; (5) through experiments and computational modeling (built with, to our knowledge, the only multicompartmental CA1 model that could generate proper synaptically driven Ca^2+^ plateaus), we continued to demonstrate that neuromodulation (acetylcholine), NMDAR and TRPM channels can shape, implement and update the BTSP state variables by affecting the strength and dynamics of plateaus.

Altogether, we advance the field by pursuing a rational deconstruction of *in-vivo* hippocampal plasticity for its place code (**Figure S12A**) in terms of building blocks for the cellular algorithms of BTSP (**Figure S12B**), and identified their biophysical underpinnings. There are two perspectives worth further discussion. To begin with, our study provides a sophisticated demonstration and consolidates the notion that dendritically initiated spikes, specifically relevant to LTP induction, are not a simple voltage-thresholded phenomenon. Rather, dendritic plateau potentials may be a substantiation of the synaptic internal states for BTSP induction. This internal state reflects the states of their inputs: electrogenesis underlying plateaus acts as an input-dynamics comparator, in such a sense that it represents a dendritic state by “listening” to the states of upstream neurons—such as activation of spatial representation sequences from CA3 recurrency (which could result in ramping firing at CA3→CA1 synapses) and activation of less spatial, phase-coupled correlates of environment information and goals in the EC/CA3 circuits (**Figure S12B**). Plateau-potential initiation cares not only about the number of co-active synapses or the total level of depolarizations, but also the timing of inputs and their dynamics, which reflects the functional states they represent. Remarkably, the dendrites enter plateau (and BTSP) states only after a sufficient amount of time has been spent on input comparison across a few theta cycles (**Figures 3B**, **4**); it is not hard to imagine that maybe the synaptic state takes several theta-cycles of time to migrate deeply enough into attractor-like rings for plateauing, in terms of nonlinear dynamics landscape. This buffering “depth” for plateauing is potentially advantageous, as BTSP is a plasticity event of ultra-high learning rates and requires tight regulations; well regulated plateaus and BTSP states can maximize low-interference yet high-capacity hippocampal learning (**Figure 7**). Dendritic spikes as a variable, graded activity is not a new idea^32,59^; however, to our best understanding, this is the first study showing that the presumably continuous synaptic states represented by dendritic spiking underlie powerful LTP—in this case, a hippocampal incarnation—that links biophysics, algorithms and *in-vivo* functional selectivity.

This work also modified the standard framework for induction of BTSP potentiation^20,31^. A most critical difference in our view is: plateau potential initiation can interact with the EPSPs to be potentiated. Besides the idea that BTSP states may be non-discrete, we also showed variation of ET dynamics as a function of activation frequency (**Figures 2**, **S3**). Thus, shifting from a simple, monolithic “ET-IS” non-Hebbian framework, in which presynaptic activation lacks frequency-dependent dynamics, this arising picture elaborates a formulation in which continuously graded presynaptic states and continuously graded postsynaptic states co-dictate the induction of BTSP. Of note, in this new paradigm, plateau potentials could serve two compatible roles: reflecting states for BTSP induction regardless of whether the input of interest is part of plateau generation, as well as offering a signal being “instructive” like an external *teacher* if the input of interest (*student*) is independent of plateau generation, possibly so by being temporally remote (> 1-2 seconds away from the plateaus)—that is to say, the notion of plateaus as an “instructor” becomes obscure when the teacher is also affected by the student. Thus, the construct of “instructive role” could be a special function under the broader concept of plateaus as *BTSP states*. We therefore suggest that plateaus as a synaptic internal state may be a more fundamental first principle, which could accommodate different views. This perspective is hard to achieve with a potentially monolithic interpretation under a simple “ET-IS” formulism of BTSP.

Our work may further provide a few provocative connections between cholinergic signaling, plasticity rules, and hippocampal algorithms and functions. Acetylcholine is a neuromodulatory system considered to mediate novelty-based synaptic learning^26^, routing of synaptic transmissions^60^ and cognitive and memory abilities^61^. These various aspects could potentially be integrated by a mechanistic explanation of Ach effects on high-fidelity synaptic strengthening we found. Recently, a proof of causal roles for MS-mediated theta-activity in hippocampal learning was provided^47^.

Moreover, theta sequences arose as a data-efficient theoretical algorithm that enabled rapid learning of observation sequences from the environment^62^. As theta rhythmicity has been considered a state of network operation that repetitively cycles between retrieval and prediction (retrospective and prospective)^45,63,64^, our finding fills a gap to address how alternating associative and predictive memory state could give a permission for potential powerful recruitment of CA1 neurons into an output of CA3 theta-organized sequences in a learning event. Although we performed experiments by compensating the reduction of CA3 input^60^ in pharmacology/optogenetics to emphasize the postsynaptic action on plateaus, acetylcholine can also bias dendritic input dynamics to prioritize the EC input. In this way, additional drive from the cortical feedback^21^ can be part of the Ach-mediated plateau facilitation. As the cholinergic functions decline in the aged brain^61,65^, our mechanistic insight may support causal understanding about the associated deterioration in low-interference hippocampal predictive encoding and memory.

That being said, we acknowledge that all of the mechanistic work done here was carried out in acute hippocampal slices. This is not *in vivo*, and we simply speculated based on principles of biophysics, neurophysiology and neural algorithms. We still do not know the nature of the signals which *inform* initiation of plateauing and the associated BTSP-LTP. As importantly, very little is known about the algorithmic nature of the presynaptic inputs, postsynaptic integration, and legitimate interactions of these factors for effective hippocampal learning. In a paper co-submitted with this study (Wang, Chen et al.), we are going to investigate these issues, which will complement the findings and interpretations of this paper. A lot more work of *in-vivo* validation and hypothesis testing can be conducted.

## STAR Methods

### KEY RESOURCES TABLE

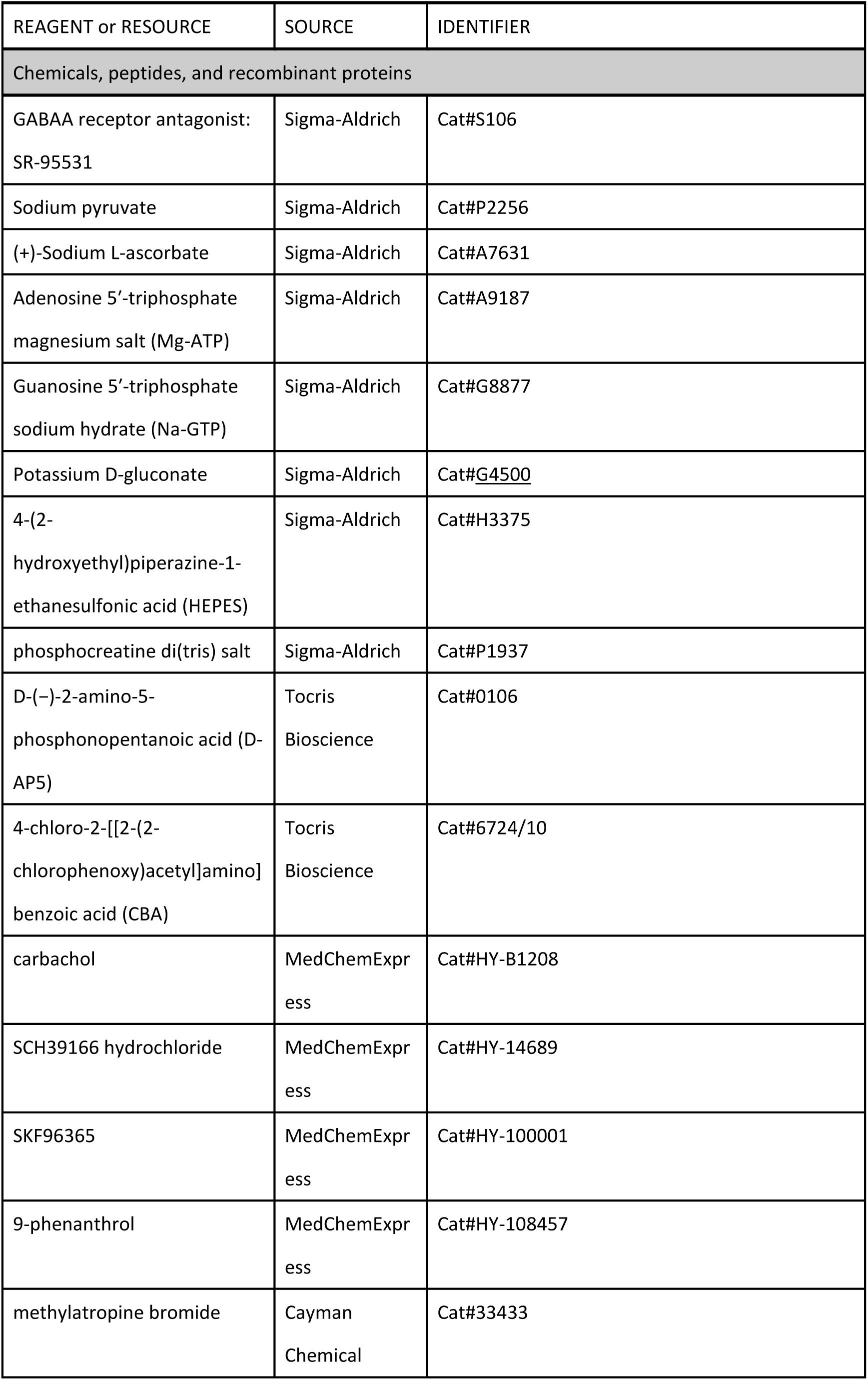
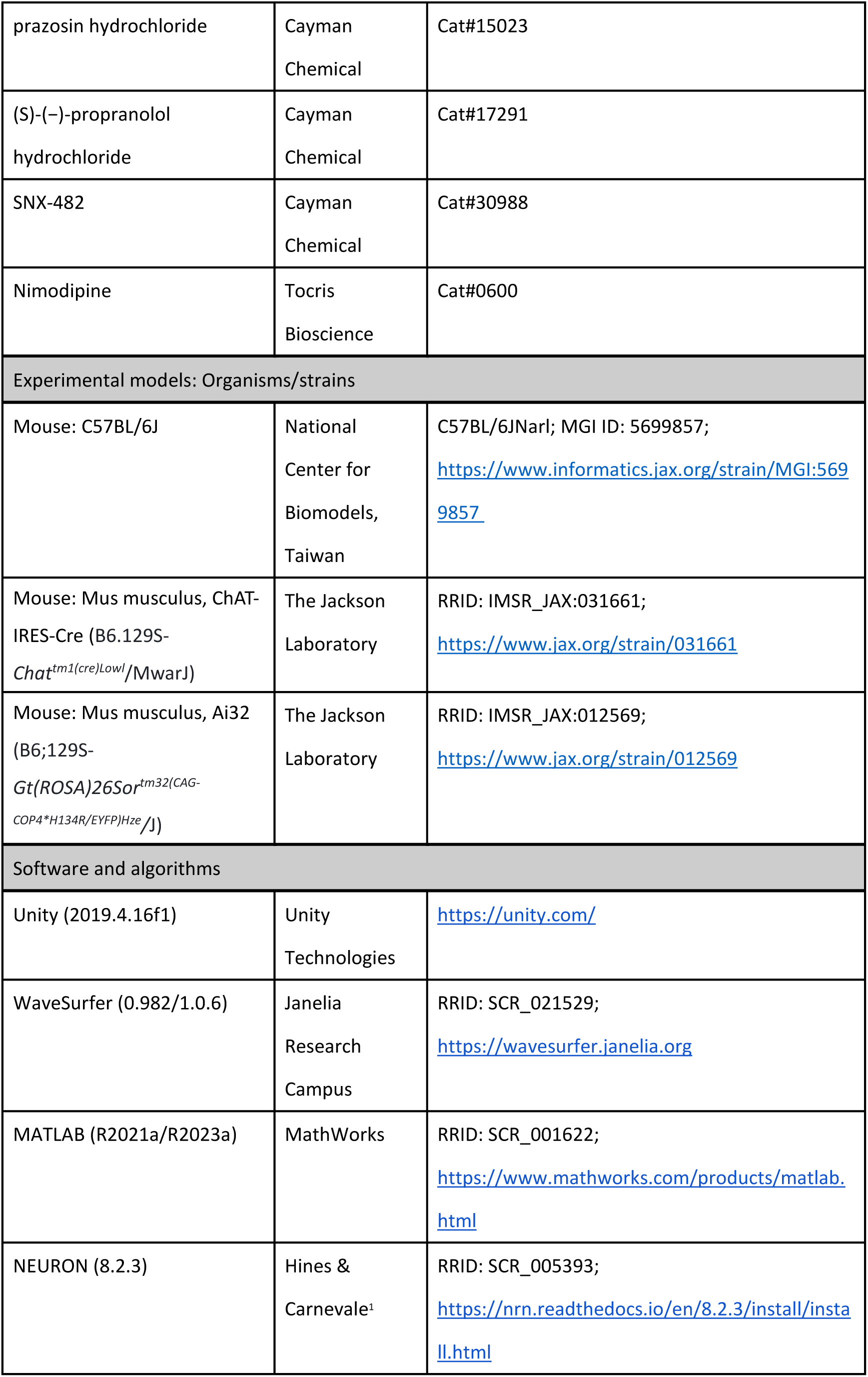
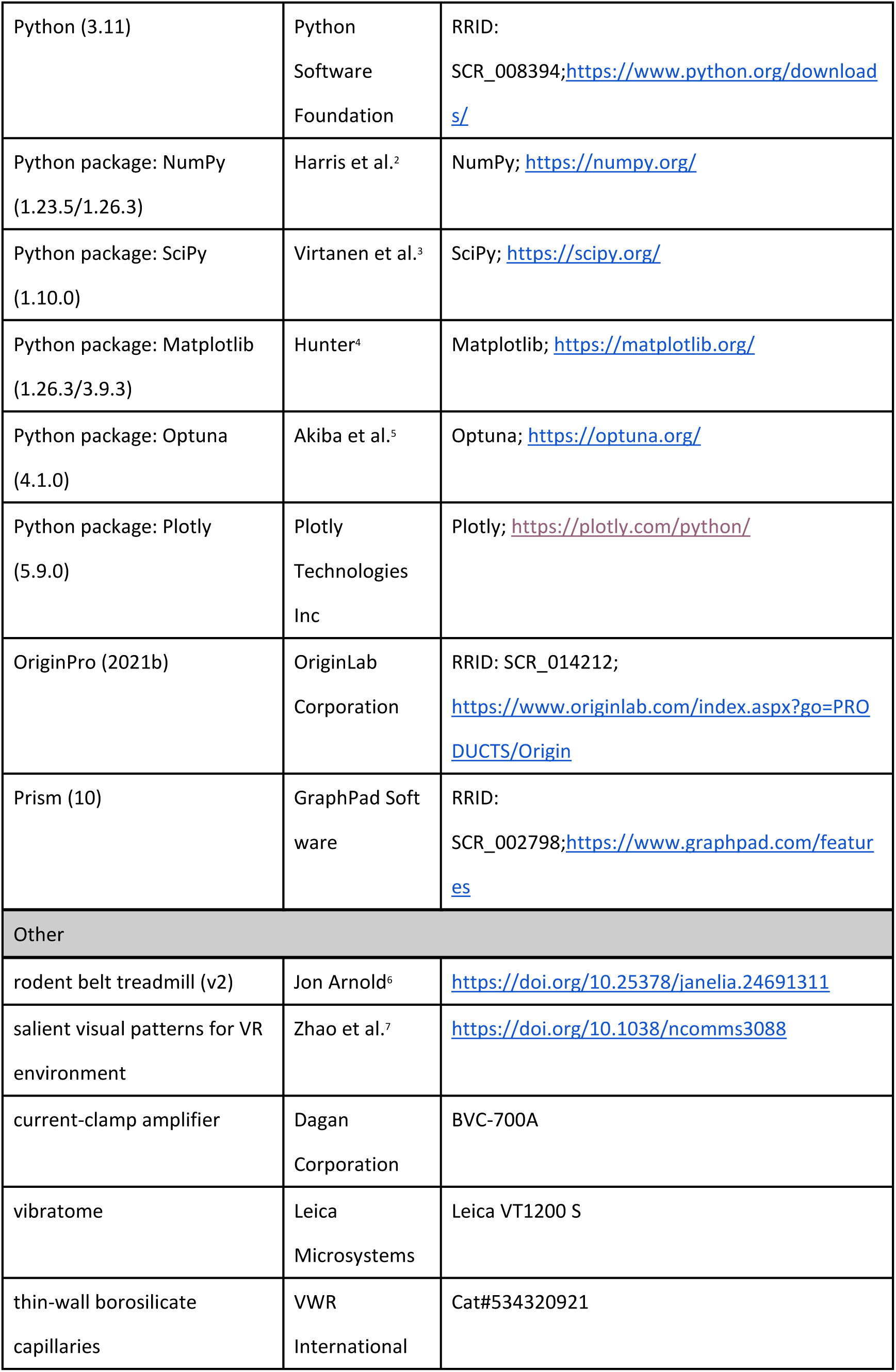

## EXPERIMENTAL AND SUBJECT DETAILS

### Mice

The use of animals in this study was approved by the Institutional Animal Care and Use Committee of Academia Sinica, and all procedures were conducted in accordance with institutional guidelines.

Male C57BL/6 mice aged 8−16 weeks were used in *in-vivo* experiment. In a 1-week recovery period after headplate surgery, daily water intake was restricted to 1 mL. During behavioral training period, animals were provided supplemental water to reach a total daily intake of 1.5 mL after each training session. Health status was regularly monitored by body weight and quantitative assessments. No blinding was employed for experiments.

Two groups of mice were used in *ex-vivo* brain slice experiments: (1) male C57BL/6N (8–12 weeks), and (2) male transgenic Ai32 and ChAT-Cre (20–25 weeks, for optogenetic experiments).

## METHOD DETAILS

### *In-vivo* experiments

#### Animal surgeries

8- to 16-week-old male C57BL/6 mice were used for *in-vivo* patch-clamp recordings. Before the onset of behavioral training, animals were implanted with a custom-made titanium head plate. The plate was secured to the skull with Zhao et al., (2013) dental adhesive resin cement (Super-Bond; SUN MEDICAL, Shiga, Japan) under isoflurane anesthesia. To minimize the number of animals used, the head plates were designed to permit bilateral access to the hippocampus. Craniotomy coordinates for electrophysiological recordings were pre-marked (relative to bregma: 2.2 mm AP, 1.7 mm ML for whole-cell recordings, and 1.5 mm posterior to this site for field recordings). Analgesia was provided with buprenorphine (0.0021 mg per 20 g body weight) at the conclusion of surgery, and ketoprofen (0.01 mg per 20 g body weight) was administered once daily for two consecutive days postoperatively.

Prior to the first electrophysiological session in a given hemisphere, craniotomies with diameters < 300 μm were performed under isoflurane anesthesia. We preserved the dura to improve mechanical stability of underlying tissues. The exposed skull and cortical surface were sealed with a silicone elastomer (KWIK-CAST; World Precision Instruments, Sarasota, FL, USA) during recovery periods and between subsequent recording sessions.

#### Mice behavioral training in virtual reality

The behavioral setup constructed in a virtual reality system was built with Unity (Unity 2019.4.16f1; Unity Technologies, San Francisco, CA, USA). Animals were head-fixed in stainless-steel holders positioned above a low-friction belt treadmill equipped with an encoder for rotation tracking (Arnold, Jon (2023). Rodent Belt Treadmill. Janelia Research Campus. Physical object. https://doi.org/10.25378/janelia.24691311.v2). The treadmill was surrounded by three thin-bezel monitors (specifications: frame rate 60 Hz), providing a visual field of ∼140° horizontally and 60° vertically from the animal’s perspective. Behavioral variables as position, speed, licking events, and rewards were all sampled at 50 Hz. The virtual environment was designed as a linear corridor measuring 342.8 cm in length and 5.7 cm in width, partitioned into 12 segments featuring salient visual patterns. These visual patterns consisted of black-and-white stripes or dot textures displayed against high-contrast grayscale backgrounds, optimized based on known tuning properties of mouse visual cortical neurons e.g., Zhao et al., (2013). Animals were trained to run unidirectionally to obtain a 10% sucrose solution reward delivered via a lick port at the middle of the 6^th^ segment (∼157 cm) of the tunnel. The later 6 segments were designed to be identical to the first 6 segments in order to create a loop-like environment. Once the mouse either successfully obtained the reward or missed the reward zone, the screen turned green for 4 seconds. At the end of the green screen , the mouse was teleported back to the starting point (0 cm) of the environment.

Beginning on the fifth day after head-plate implantation, mice were provided with a saucer running wheel in the home cage. Following a one-week recovery period, they were placed on a water-restriction protocol (1 mL water/day). Behavioral training began after one week of restriction. After a brief acclimation session to the setup, animals underwent ∼20 days of daily training, with session length progressively increased to 60–70 min. Following each session, supplemental water was given to maintain a total daily fluid intake of 1.5 mL.

During typical recording sessions conducted after training, mice consistently engaged in continuous running without observable signs of distress or discomfort. They also displayed anticipatory licking upon approaching reward sites, indicative of learning the spatial contingencies of the task.

#### In-vivo whole-cell patch-clamp recording

Whole-cell patch-clamp recordings were performed 0.5–1 day after craniotomy and continued for 2–3 consecutive days in each hemisphere. To determine the depth of the somatic layer of pyramidal cells (stratum pyramidale) within dorsal hippocampal CA1, we first inserted a long-taper extracellular pipette (electric resistance ∼1 MΩ when filled with 0.9% NaCl) vertically through the rostral craniotomy. Theta-modulated spiking and sharp-wave activity were detected at depths of ∼1.1–1.3 mm below the brain surface. After determining the depth, a glass pipette for local field potential recordings was advanced obliquely (45°) through the caudal craniotomy until positioned near CA1 stratum pyramidale, guided by trigonometric calculation. The pipette was then discarded, and a long-taper patch pipette (electric resistance 8– 13 MΩ, filled with the same K-gluconate-base intracellular solution used for slice electrophysiology) was introduced vertically into the rostral craniotomy. Positive pressure of 10–11 psi was applied while traversing neocortex and reduced to ∼0.4 psi upon entry into hippocampal tissue. Somatic whole-cell access was obtained using the “blind-patch technique” (Blanton et al., 1989). Recordings were accepted only when exhibiting canonical CA1 pyramidal cell signatures (Harvey et al., 2009; Lee et al., 2012). After break-in, the characteristic activity of CA1 pyramidal neurons often displayed a brief quiescent period with hyperpolarized membrane potentials (*V*_m_) lasting <1 min, occasionally up to ∼3 min, before stable subthreshold fluctuations and theta oscillations emerged. Analyses were restricted to periods where *V*_m_ was stable, typically −70 to −50 mV. Patch-pipette series resistance was typically ranged from 20– 80 MΩ and was monitored throughout the experiment. Bridge balance was carefully monitored and adjusted during all recordings.

Of 19 neurons recorded with standard intracellular solution, 11 exhibited spatially modulated ramp-like synaptic responses that drove action potential firings after induction, while the others remained silent. The spatially tuned fields were induced with brief, large somatic current injections (600 pA, 300 ms) (Bittner et al., 2015) to test voltage dependence of synaptic responses.

Current-clamp recordings were performed with a Dagan BVC-700A amplifier (Dagan Corporation, Minneapolis, MN, USA). Data were filtered at 5 kHz (low-pass) and digitized at 20 kHz using a BNC-2110 interface (National Instruments, Austin, TX, USA). WaveSurfer software (1.0.6; Janelia Research Campus, Ashburn, VA, USA) was used for data acquisition.

### *Ex-vivo* experiments

#### Acute hippocampal brain slice preparation

Male C57BL/6N mice (8–12 weeks) and male transgenic ChAT-Cre/Ai32 mice (20–25 weeks, used for optogenetic experiments) were deeply anesthetized with isoflurane and subsequently decapitated. Brains were rapidly extracted and placed in ice-cold dissection solution containing (in mM): 204.5 sucrose, 2.5 KCl, 1.25 NaH_2_PO_4_, 28 NaHCO_3_, 7 dextrose, 3 Na-pyruvate, 1 Na-ascorbate, 7 MgCl_2_, and 0.5 CaCl_2_ (pH 7.3, 310–320 mOsm, oxygenated with 95% O_2_ and 5% CO_2_). Hippocampal slices (350-μm thick) were cut at an oblique angle using a vibrating tissue slicer (VT1200S; Leica Microsystems, Wetzlar, Germany). To prevent polysynaptic recruitment during synaptic stimulation, the CA3 region and superficial layers of the entorhinal cortex were removed.

Slices were transferred onto a suspended mesh in an incubation chamber filled with artificial cerebrospinal fluid (ACSF) composed of (in mM): 125 NaCl, 2.5 KCl, 1.25 NaH_2_PO_4_, 25 NaHCO_3_, 25 dextrose, 2 CaCl_2_, 1 MgCl_2_, 3 Na-pyruvate, and 1 Na-ascorbate (pH 7.3, 295–300 mOsm; continuously bubbled with 95% O_2_ and 5% CO_2_). Slices were recovered for 30–60 min at 35–37 °C, after which the chamber was maintained at room temperature.

#### Ex-vivo whole-cell patch-clamp recording

Recordings were performed in submerged slices placed in the recording chamber of an upright microscope (BX51WI; Olympus, Tokyo, Japan) equipped with infrared differential interference contrast (IR-DIC) optics and a water-immersion objective (40 x, 0.80 NA or 60x, 1.00 NA; Olympus, Tokyo, Japan). Slices were continuously perfused with oxygenated ACSF maintained at 31–33 °C. The GABA_A_ receptor antagonist SR-95531 (1–2 µM) was routinely applied.

Patch electrodes were pulled from thick-wall borosilicate glass to yield electric resistances of 6–8 MΩ when filled with intracellular solution. For multiple pathway stimulation experiments, the internal contained (in mM): 130 K-gluconate, 10 KCl, 10 Na22-phosphocreatine, 10 HEPES, 4 Mg-ATP, 0.3 Na-GTP (pH 7.3, 295–300 mOsm). For experiments characterizing frequency-dependent eligibility traces and regenerative Ca^2+^ spiking plateaus, the internal contained (in mM): 130 Cs-methanesulphonate, 6 KCl, 10 HEPES, 4 NaCl, 2 Mg-ATP, 0.3 Na-GTP, 14 Tris-phosphocreatine (pH 7.3, 290–300 mOsm). Reported membrane potentials were not corrected for liquid junction potential. For experiments examining dendritic plateaus with or without regenerative activity, long-taper electrodes were fabricated from thin-wall borosilicate capillaries (534320921; VWR International, Radnor, PA, USA) and had electric resistances of 4–8 MΩ.

Whole-cell current-clamp recordings were obtained using a Dagan BVC-700A amplifier. Signals were low-pass filtered at 5 kHz and digitized at 50 kHz with National Instruments BNC-2090 and BNC-2110 boards. Acquisition was controlled by WaveSurfer 0.982/1.0.6 running in MATLAB (MathWorks, Natick, MA, USA).

Stimulating electrodes were placed in the stratum lacunosum-moleculare (SLM) ∼400–500 μm and the stratum radiatum (SR) ∼200–300 μm away (toward subiculum) from the recorded cell to stimulate the perforant path (PP, also known as temporoammonic path) and the Schaffer collaterals (CA3_2_), thus recruiting synaptic inputs from the EC and the CA3 region, respectively. For generation of synaptic driven dendritic plateau potential (Takahashi & Magee, 2009), theta-burst stimulation (TBS) which consisted of five high-frequency bursts (5 pulses at 100 Hz) repeated at an inter-burst interval of 200 ms (5 Hz) was simultaneously delivered at PP and CA3_2_ pathway. A stimulating electrode was also placed in the SR (CA3_1_) to evoke ramp-like EPSPs as spatially tuned subthreshold synaptic inputs onto a recorded CA1 pyramidal neuron. The ramp-like stimulus patterns were symmetric and the stimulus used was generated with a peak frequency (at the center of the ramp) of ∼50 Hz with a total duration of 1.2 s (Hsu et al., 2018). Five pairings of TBS (PP + CA3_2_) with a ramp-like stimulus (CA3_1_) were delivered either synchronously or in separate time windows to induce long-term potentiation (LTP). Initial amplitude of EPSPs recorded from PP, CA3_1_ and CA3_2_ pathways were set to ∼5 mV. Pathway independence was verified using paired-pulse ratio (PPR) analysis before recordings. To confirm non-overlapping CA3_1_ and CA3_2_ afferent, the two pathways were alternately stimulated with a paired-pulse protocol (50 ms interval), and PPR was calculated as EPSP_2_/EPSP_1_. Both CA3_1_ and CA3_2_ exhibited paired-pulse facilitation (PPR ≈ 2), whereas cross-stimulation (single CA3_1_ pulse and followed by single CA3_2_ pulse) yielded a PPR near 1. One-way ANOVA (*n=*15) revealed no difference between CA3_1_ and CA3_2_ (*n.s.*), but both were significantly higher than cross-stimulation (*\*\*\*p* < 0.001) (data not shown). In the experiment of frequency-dependent eligibility trace, CA3 input stimulation containing 10 pulses with various frequencies was paired with a CA1 somatic current injection (300 ms, 600 pA) to induce dendritic plateau potential through cesium-base internal solution (Bittner et al., 2017). This pairing protocol was delivered either simultaneously or separated in different intervals.

Experiments for with and without regenerative spiking plateau potentials used the same protocol, pairing 20 Hz CA3 input with postsynaptic current injection at Δ*t* = 0. Paired-pulse EPSPs (50 ms interval) were monitored every 45 s to assess synaptic strength, and interleaved test pulses were used to monitor the bridge balance, pipette capacitance compensation, and input resistance of the cell throughout the experiment.

#### Pharmacology

Chemicals were bath-applied for 10-12 minutes before subsequent experiments commenced. A typical experiment (**Figures 6, 9, S10**) started with a ∼10 minutes recording as the baseline condition, followed by 7-8 minutes of a pharmacological agent applied in the bath solution. Next, if a depression of EPSPs was observed, the stimulus intensity was recalibrated to compensate for the loss of synaptic excitation at the dendrites, and then, upon 10-12 minutes total of drug application, LTP induction protocol was applied, followed by ∼8 minutes washout before the regular monitoring of EPSP was resumed.

A big body of literature has applied Carbachol in the bath solution to examine the functions of muscarinic Ach signaling in slice experiments. We were full aware of the caveat of using an agonist at a constant concentration over time, which could lead to different results compared to transient activation of the signals with optogenetics (Rosen et al., 2015). Our key conclusions were not made without an argument based on antagonism (in this case, Atropine), which supports the effects of a given signaling from endogenous sources. Nevertheless, as far as the use of an agonist is concerned (**Figure 6**), careful literature review as well as control experiments with Carbachol washed in and then washed out (with no LTP induction) indicate the following, particularly about any effects at around 20-30 minutes after the induction (when we evaluated synaptic plasticity): (a) with CCH at relatively lower concentrations (< 3-5 uM), longer application (> 15 mins) led to an LTP (399 ± 109 %, n = 3 cells; range = 289—616 %), consistent with previous work (Auerbach & Segal, 1996; Yun et al., 2000); (b) with CCH at a relatively higher concentration (> 10 uM), longer application (> 15-20 mins) led to significant LTD (Benoy et al., 2021; Dickinson et al., 2009; Hasselmo & Schnell, 1994); (c) with CCH at several concentrations (2—10 uM), shorter applications (< 15 mins) did not result in long-lasting effects of EPSP amplitude change at the time following the wash-out (**Figure 6B**; 150 ± 17 %, n = 4 cells). Lastly, in our initial experiments, ∼45 mins of CCH bath-application with plateau induction led to a continual growth of EPSP up to 400-500% at 45 mins after induction (n = 3 cells), which led to speculation that a prolonged CCH condition would result in long-lasting effects. In conclusion, these cautious considerations led us to adopt a concentration of 10 uM CCH, an effective dose used in the literature (e.g., Palacios-Filardo et al., 2021), but being applied for only 10-12 minutes to avoid any undesired long-term effects of CCH bath-application on the CA3→CA1 synaptic transmission exhibited at the time window we measured LTP.

Reagents were obtained from the following vendors: Sigma-Aldrich (St. Louis, MO, USA): SR-95531, sodium pyruvate, (+)-sodium L-ascorbate, phosphocreatine disodium, magnesium adenosine 5′-triphosphate (Mg-ATP), sodium guanosine 5′-triphosphate (Na-GTP), 4-(2-hydroxyethyl)piperazine-1-ethanesulfonic acid (HEPES), potassium D-gluconate, and dextrose; Merck KGaA (Darmstadt, Germany): calcium chloride, magnesium chloride, sodium chloride, potassium chloride, sodium bicarbonate, and sucrose; Tocris Bioscience (Minneapolis, MN, USA): D-(−)-2-amino-5-phosphonopentanoic acid (AP5), nimodipine, and 4-chloro-2-[[2-(2-chlorophenoxy)acetyl]amino]benzoic acid (CBA); MedChemExpress (Monmouth Junction, NJ, USA): carbachol, SCH39166 hydrochloride, SKF96365, and 9-phenanthrol; Cayman Chemical (Ann Arbor, MI, USA): methylatropine bromide, prazosin hydrochloride, (S)-(−)-propranolol hydrochloride, and SNX-482. No blinding was applied in these experiments.

#### Optogenetics

All experiments were conducted using male C57BL/6J mice as the background strain. Cre reporter allele mice (Chat-IRIS-Cre) were used to label cholinergic neurons. Homozygous Chat-IRIS-Cre mice were crossed with homozygous Ai32 mice, resulting in heterozygous offspring expressing channelrhodopsin-2 (ChR2).

A pE-300 light engine (CoolLED, Andover, UK) was used for optogenetically evoking the endogenous release of acetylcholine from cholinergic inputs in the hippocampal slices. Blue light (460 nm, 5 ms, ∼5mW/mm^2^) was delivered to slices at 5 Hz for 5 min through the 40x microscope objective.

#### Computational modeling for synaptic plasticity

We performed BTSP kernel simulations in Python (3.11; Python Software Foundation, Wilmington, DE, USA) using NumPy (1.23.5) and SciPy (1.10.0), optimized parameters with Optuna (4.1.0), and plotted results using Matplotlib (3.9.3).

#### Data fitting using the Milstein-Romani model

The BTSP model formulation and a subset of parameters were adapted from Milstein et al., (2021) with modifications. We considered only the LTP component, and synaptic weight change was calculated as:

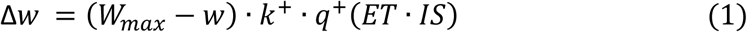

where Δ*w* was weight change, *w* was current weight, *W_max_* was maximum attainable weight, *k*^+^ was learning rate, and *q*^+^ was sigmoidal function of the product of two signals: eligibility trace (ET) and instructive signal (IS). The *q*^+^ was defined by the generalized sigmoidal function *s*(*x*, *⍺*, *β*), where *⍺* and *β* could control the offset and slope.

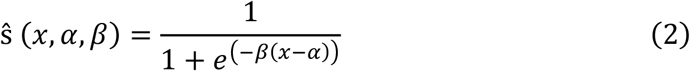

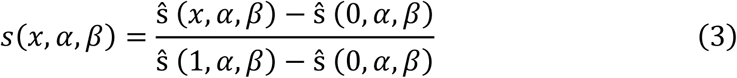

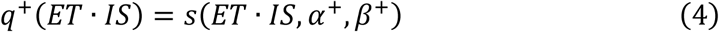

The ET was modeled as:

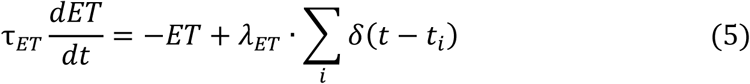

where τ*_ET_* was the time constant, *λ_ET_* was the scaling factor, *d*_*i*_ represented the times where pulse inputs were given. To simulate experimental conditions, we used the following input patterns: fixed-rate inputs (10, 20, 40, 100 Hz) of 10 pulses. The plateau-induced instructive signal (IS) was modeled as:

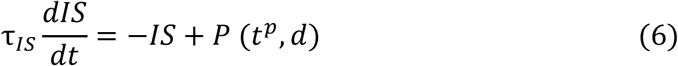

where τ*_IS_* was the time constant, *P* was a binary function where the outcome was 1 during plateau and 0 otherwise, *d*^*p*^ was the plateau induction timing, and *d* was the plateau duration. It should be noted that after we obtained the IS, we performed the peak normalization to 1 and multiplied the IS by the scaling factor.

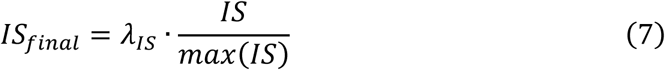

To derive the BTSP kernel, we fixed IS timing at 0 (*d*^*p*^ = 0) while sliding the input from −5 to +5 s, where weight changes approached zero at both extremes. From this kernel, we calculated the peak amplitude of BTSP kernel and the backward time constant.

During parameter optimization. We only allowed two parameters, τ*_ET_* and *λ_ET_*, to be optimized while fixing other parameters. We optimized these parameters using the Python package Optuna to match two target values observed: peak amplitude of BTSP kernel and the backward time constant, for each experimental condition.

#### Modeling of BTSP kernels with different forms of IS

Based on Milstein-Romani model, we created three different variations of IS generation in these BTSP models. The overall BTSP model followed the same structure as Eqs. (1)–(5), except that here the input pattern is “ramp”, and IS (Eq. (6)) underwent further modulation according to different assumptions.

Cond. (i): IS from plateau with graded duration and graded scaling factor. First, plateau duration was determined by a Gaussian function that peaks at Δ*t* = 0 between the input stimulus and plateau. This modulated duration became a parameter for Eq. (6). Under cond. (i), there was no normalization as mentioned in the section “Fitting of the Milstein-Romani model”, Eq. (7).

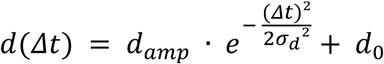

Second, IS derived from Eq. (6) was multiplied by another scaling factor determined by a Gaussian function, which also peaked at Δ*t* = 0 s between the input stimulus and plateau.

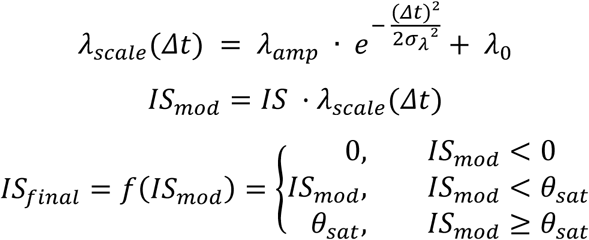

Third, the modulated IS underwent a nonlinear piecewise function. The final IS was then applied to Eq. (1) to obtain the synaptic weight change.

Cond. (ii): IS from plateau with two-level (low and high) scaling. The plateau duration modulation was the same as in cond. (i). However, in cond. (ii), the normalization existed as stated in the section “Fitting of the Milstein-Romani model”, Eq. (7). This meant that duration had little effect. Next, IS was scaled depending on whether Δ*t* fell within the ±0.5 s window or not. This scaling factor was the maximum value of Gaussian function from cond. (i). Finally, the modulated IS underwent a nonlinear piecewise function and was applied to Eq. (1), outputting the synaptic weight change, same as cond. (i).

Cond. (iii): IS from plateau with two-level (low and high) duration modulation and scaling factor, and had a definitive cutoff. This condition was the same as cond. (ii), but with a cutoff at Δ*t* = ±0.5 s. Outside the cutoff value, there was no plateau, and therefore no IS.

#### Modeling neuromodulator-modulated LTP and hippocampal encoding capacity

We modeled the CA1-CA3 region as a network of rate units. There was an input layer corresponding to CA3 cells, a hidden layer corresponding to CA1 and a tacit readout layer. The input units modeled a context-dependent place code in CA3. The rate of each unit x = [*x*_1_*, x*_2_*, …, x*_N_ ] depended on the position of the agent in discretized space with bin size Δ*X*, such that the position *X* corresponded to an index *k*, *X* = *k*Δ*X*. CA3 place code was modeled by an indicator function (Kronecker delta) *x_i_* = *δ_kl_* such that *x_i_* = 0 if *X* =*̸ l* and *x_i_* = 1 otherwise. In this way, the size of the CA3 place field was the size of the spatial bins. Each CA3 unit was assumed to code for a single position in a one-to-one fashion, such that a given environment had a fixed one-to-one lookup table between unit indices *i* and position indices *k*. We denoted the index mapping function for environment *m* as *k* = *h_m_*(*i*). Each new environment was modeled by scrambling the one-to-one lookup table. Once created, the lookup table was associated with the new environment such that the agent can return to a previously seen environment by using the lookup table for that environment.

Each input unit mapped to a single CA1 unit with weight vector *w* = [*w*_1_*, w*_2_*, …, w_N_* ] such that at each moment in time, the input current was given by *I*(*X*) = **w***^T^* **x**(*X*), where *X* was the current location. The positions were visited sequentially in time with a constant speed, *v*, such that *X* = *i*vΔ*t* for time index *i* with time bin Δ*t*. Since we used an indicator function for the CA3 place code that matched the spatial bins, this operation implied that at each moment in time, the input to the CA1 unit was the weight of the connection from the CA3 unit (index *k*) that was active at that time *I_i_* = *w_k_* = *w_hm_*_(*i*)_. Thus, in this model, a given environment sequentially samples from the same weight vector, with different environments modeled with different sampling order. The output of the CA1 cell was a nonlinear readout of the potential *r_CA_*_1_ = *f*(*V_i_*), where the potential was modeled as a simple temporal smoothing over *T* time bins of the input current *V_i_* =^1^*_T_*P*^i^_j_*_=*i−T*_*I_j_* .

The weight vector was initialized by randomly sampling a Gaussian distribution. Simulated environment crossings altered the weights through plasticity. Plasticity was modeled as unfolding according to a supervisory signal in the apical dendrites of CA1, itself inherited in part from the entorhinal cortex (Grienberger & Magee, 2022). A plasticity rule altered the weights towards the supervised signal *t_i_*, chosen to be a Gaussian centered at a target location *x*_tar_. We assumed that the target was subtracted from the activity of the cell in the apical dendrite such that the potential there was *t_i_−f*(*V_i_*). Plateaus could be generated only when this difference was positive, hence we required a factor floor *t_i_ − f*(*V_i_*) *−* Θ*_E_⌋* for the generation of a plateau, where Θ*_E_* was a parameter regulating the size of the error between supervision and activity that engaged the plateau. We must further included an additional factor to take into account that complex CA1 spikes occurred with a conjunction of basal inputs from the CA3 and apical inputs from EC (Bittner et al., 2015; Takahashi & Magee, 2009). To further take into account that the relationship between CA3 inputs and plateau probability was non-monotonic (Park et al., 2025), we considered the non-monotonic function to be the derivative of the activation function *f^ʹ^*(*V_i_*). We modeled the plateau-duration dependence of LTP by a linear scaling of the plateau duration *T_P_* . Lastly, the plasticity rule was modeled as a Hebbian conjunction of a second-long presynaptic eligibility trace *E_i_* and the postsynaptic plateau, together we had four factors multiplied, namely the plateau duration, the cell-body activation, the apical dendrite activation and the pre-synaptic eligibility trace:

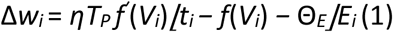

Depression was modeled by ensuring a normalization of the weight vector at every timestep, an operation that could arise from the saturation of individual synaptic elements (Madar et al., 2025; Milstein et al., 2021; Oja, 2008) and dendritic organization of synapses (Bird et al., 2021).

We modeled a novelty signal as being provided by NE or ACh inputs to the hippocampus. This novelty signal was set as high when a new environment occured and gradually decreased with repeated exposures to the same environment according to an exponential function of time. The effect of this novelty signal was to increase the plateau duration *T_P_* , such that *T_P_* = (*e^−i/τ^* + *b*)*/*(1 + *b*). The agent was modeled as being exposed to one environment repetitively, then switching to a new environment after a number of exposures L = 5000.

A decoded position was extracted from the firing of the CA1. We used moment of the activity through one environment crossing as the decoded position. We quantified storage error as the difference between the decoded position and the target position after the L exposures to the environment. We quantified recall error as the difference between the decoded position and the target position when returning to the first environment after exposure to one or many environments.

### Multicompartmental CA1 pyramidal neuron modeling

Simulations were performed with NEURON 8.2.3 (Hines & Carnevale, 1997) via Python interface (3.11). Python packages, including NumPy (1.26.3), Matplotlib (1.26.3), and Plotly (5.9.0) were used to facilitate stimulation protocol implementation, data analysis, and data visualization.

#### Parameter tuning

The morphological reconstruction and synaptic distribution of the simulated hippocampal CA1 pyramidal cell were based on the previous work of Bloss and Cembrowski (Bloss et al., 2018). Parameters for passive properties were kept identical as in their model. For our proposes in studying calcium-related electrophysiology, we deployed additional mechanisms into the model on top of those originally included in Bloss and Cembrowski. The newly introduced mechanisms included: Ca^2+^-holding mechanisms, L-type Ca^2+^ channel, R-type Ca^2+^ channel, slow-AHP type Ca^2+^-dependent K^+^ channel, medium-AHP type Ca^2+^-dependent K^+^ channel, TRPM channel, and HCN channel. The kinetic profiles of Ca^2+^-holding mechanisms, Ca^2+^ channels, Ca^2+^-dependent K+ channels, and HCN channel were from Poirazi et al. (2003) The kinetic profile of TPRM channel was from (Combe et al., 2023). For simplicity, it was assumed that cholinergic modulation of TRPM channel was always present in this study.

The active properties (both those that had been present in Bloss-Cembrowski model and those introduced by us) were tuned altogether in order to meet three major constraints: (1) the linear and supralinear transition of somatic EPSP elevation with increasing numbers of stimulated synapses in single-branch glutamate uncaging experiments (Losonczy & Magee, 2006), (2) the formation of complex spikes in dendritic current injection, which was prevented by Ca^2+^ channel blocker (Golding et al., 1999), and (3) the generation of dendritic plateaus after receiving simultaneous theta burst inputs from CA3 and EC pathways for a sufficiently long time (>600 ms) (Takahashi & Magee, 2009). The mean conductance and distribution of the ion channels in Bloss & Cembrowski or Poirazi & Mel were used as the starting point of our tuning. Due to the limitation of manual tuning, the tuning process was carried out in a semi-independent manner between constraints. The mean conductance of A-type K^+^ channel and the NMDAR channel were first tuned to meet constraint (1). Then, the mean conductance and distribution of Ca^2+^ channels, Ca^2+^-dependent K+ channels, TRPM channel, and HCN channel and the slow inactivation kinetics of Na^+^ channel were tuned to meet constraints (2) and (3). In general, a higher conductance of L-type Ca^2+^ channel increased the amplitude and duration of the bursts immediately following deliveries of stimulation; a higher conductance of R-type Ca^2+^ channel gave rise to spiking and dampened repolarization between each theta burst; a higher conductance of TRPM channel led to elevation in afterdepolarization potential; a higher conductance of slow-AHP type Ca^2+^-dependent K+ channel led to a decrease in afterdepolarization potential; a higher conductance of medium-AHP type Ca^2+^-dependent K^+^ channel accelerated repolarization and prevented spiking soon after deliveries of stimulation; and a higher extent of Na^+^ channel slow inactivation prevented spiking during theta burst intervals. Tuning the parameters of each channel in an opposite way would do vice versa. However, due to the nonlinear nature of the ion channels and the even more complicated interactions between them, these were at most crude principles induced from hands-on experience in tuning and sometimes an arbitrary guess would still be required to make a better fit to constraints. After the model was tuned for constraints (2) and (3), we recursively checked if the newly-tuned parameter set still held true for constraints (1), until the parameter set was confirmed to satisfy all the aforementioned constraints.

Other properties of the modeled neuron, including input resistance at different locations, attenuation along dendritic trunk, and AMPA/NMDA ratio were also checked to be in reasonable ranges in comparison to experimental measurements. ***Synaptic stimulation*** Spatial distribution and the corresponding input source of each synapse were kept identical as the anatomical data in Bloss and Cembrowski. Spines were not explicitly modeled. Deliveries of stimulation consisted of two parts: (1) the vecevent object encoding the times of stimulation connected to a neurotransmitter release probability mechanism (2) the release probability mechanism connected to the stimulated synapses’ AMPAR and NMDAR mechanisms. Theta burst stimulation was delivered as a series of stimulus trains, each consisting of five pulses at 100 Hz (i.e., 10 ms interpulse interval). The trains were repeated at 5 Hz (i.e., with a 200 ms interval between the onset of consecutive trains). Ramp frequency stimulation was designed to represent the inputs from CA3 when animals were traversing through the place field based on previous studies. The release probability mechanism was based on the previous work of (Grienberger et al., 2017) to account for the short-term plasticity at synaptic sites. Release probability was set to be constant one for glutamate uncaging simulation. The conductance of AMPAR and NMDAR was set to be identical throughout the dendritic tree in this study. 10% of inflow from NMDAR channel was taken as calcium current and incorporated into calcium-related mechanisms. The selection of stimulated synapses was implemented by the built-in random function in Python. The number of synapses to be stimulated from each pathway was decided, and the specific synapses were selected in a non-repetitive random manner using an arbitrarily assigned random seed, with each synapse within a given pathway having a uniform probability of being selected.

### Pharmacology simulation

For pharmacological simulations, identical random seeds for synapse selection were used across control conditions and each pharmacological condition for comparability. The conductance of the ion channels simulated to be blocked was set to be 0 if not otherwise stated. For simulation of no short-term plasticity, the release probability was set to be constantly 0.5.

## QUANTIFICATION AND STATISTICAL ANALYSIS

### *Ex-vivo* experiments

The electrophysiological data were analyzed using MATLAB R2021a and OriginPro 2021b (OriginLab Corporation, Northampton, MA, USA). Statistical analyses were carried out using OriginPro 2021b, Prism 10 (GraphPad Software, San Diego, CA, USA), and Microsoft Excel (Office 2019; Microsoft Corporation, Redmond, WA, USA). All data were reported as the mean *±* standard error of the mean (SEM). Before conducting inferential statistics, the distribution of all datasets was assessed for parametric conditions using a Shapiro-Wilk test for normal distribution and an *F*-test for equal variance.

In cases of normal distribution, three-pathway induced BTSP-LTP was analyzed using repeated measures (RM) one-way analysis of variance (*ANOVA*). The contribution of NMDA receptors on the early and late components of BTSP-LTP was assessed using two-tailed unpaired Student’s *t*-tests for comparisons between control and APV treatment groups. The effects of pharmacological treatment on BTSP-LTP were analyzed using two-tailed unpaired Student’s t-tests for comparisons between two groups and independent samples. The effects of various induction protocols on BTSP-LTP were evaluated using multiple comparisons conducted by one-way *ANOVA*. When the *F*-ratio was significant, all-pairwise post hoc comparisons were made using Tukey’s post-hoc test. For the characterization of the plateau index, the *V*_m_ integral area under the 3rd to the 5th burst was normalized by the *V*_m_ integral area under the 1st burst. The relationship between BTSP-LTP and plateau index was analyzed using regression-based fitting with linear, exponential, and sigmoid models. Model adequacy was assessed using the coefficient of determination (R^2^), residual diagnostics, and hypothesis testing of the regression coefficients. Nested model comparison was performed using an *F*-test.

In the experiment of frequency-dependent eligibility trace, BTSP-LTP kernel time constants were calculated across various CA3 input frequencies, and fit with exponential regression.

To probe the effect of cesium on membrane excitability, we quantified the extent of Cs⁺ dialysis. This quantification was formulated on the basis of Fick’s law of diffusion, which describes the movement of molecules from regions of higher to lower concentration:

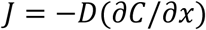

where *J* was the diffusion flux, *D* was the diffusion coefficient, *C* was the concentration, and *x* was the distance. By combining the law with expressions 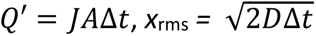, and *R*_s_ ∝ *L*/*D*, we calculated the Cs⁺ dialysis factor *Q* (s^2^/MΩ^2^) as a function of the series resistance of recordings, *R*_s_, and the duration from whole-cell access to plateau induction, Δ*t*:

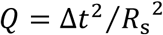

Here, *Q*^′^was the total diffusion amount, *A* was the cross-sectional area assumed as constant, *x*_rms_ was the characteristic diffusion distance (the root mean square displacement of Cs⁺), and *L* was the diffusion path length.

Our working definition of plateau activity with and without regenerative events was as follows: (1) the analysis window was the 300-ms period during which somatic current injection was paired with CA3 presynaptic input; (2) plateau traces without regenerative activity were characterized by a sustained, flat depolarization, with minimal repolarization other than the initial transient at current injection onset; (3) plateau traces with regenerative activity were defined by recurrent spiking within the plateau duration, each spike showing a peak overshoot followed by a transient drop in *V*_m_ during repolarization. For grouping, only neurons whose plateau responses across all five sweeps matched the above criteria were classified as the without-regenerative-activity subset.

We chose balanced accuracy (bACC) as the evaluation metric for selecting the optimal threshold of each predictor, *Q* and *R*s. The metric was calculated according to the equation:

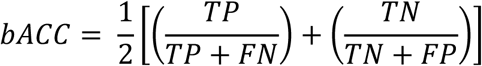

where *TP* denoted true positives, *FN* denoted false negatives, *TN* denoted true negatives, and *FP* denoted false positives. Given that bACC is particularly suited for assessing classifier performance with imbalanced datasets, it provided a fair measure between the two regenerative activity classes (with: *n* = 43; without: *n* = 14). To visualize performance, bACC was plotted as a function of thresholds, with each threshold defined at the midpoint between two adjacent data points.

### *In-vivo* experiments

#### Spatial profile of lick rate and running speed

Behavioral data were read by Arduino and output by Unity at ∼50 Hz. The time series of lick rate was calculated by differentiating time series of licks. Spatial profiles of lick rate and speed were binned by 2.86 cm.

#### In-vivo whole-cell patch-clamp recording

Electrophysiological recordings were first aligned with behavioral data using sync-pulses. Behavioral data (licks, speed, position, trial number etc.) were then interpolated to match the sampling rate of electrophysiological recordings (20 kHz).

#### Spike detection

To analyze firing rates, d*V*_m_/d*t* was calculated, and action potential onset was defined as d*V*_m_/d*t* > 5 mV/s. After spike detection, data were manually checked to minimize false positives and false negatives. In noisy recordings, action potential threshold was adjusted to d*V*_m_/d*t* > mean(d*V*_m_/d*t*) + 8*sigma to reduce false negative effects.

#### Firing-rate map

To calculate the firing rate of each position, the entire track was binned by 2 cm. The firing rate in each bin was computed as spike counts divided by occupancy time. To plot the spatial profile of the firing rate, raw traces were filtered by a Gaussian filter with 2 cm SD.

#### Subthreshold V*_m_* ramp

Spikes were first removed by discarding points 0.26 ms before and 3.4 ms after the threshold value mentioned above. The resulting trace was low pass filtered (< 3 Hz) using an FIR filter with a 200 ms Hamming window. To plot the spatial profile of the subthreshold *V*_m_, baseline *V*_m_ (defined as the average number of 20000 minimal subthreshold *V*_m_ data points corresponding to 1 second) was subtracted from the trace. After baseline subtraction, the trace was binned by 0.1 cm and smoothed by a Gaussian filter with 2.5 cm SD.

#### Theta power

To compute the theta power of each recorded cell, the *V*_m_ data around current injection were removed, and the effect of action potentials was suppressed using a median filter with 3 ms window. Since theta oscillations are notably prominent during movement, data with speed < 2 cm/s were excluded. The first 3 min of the data were used to estimate power spectral density (PSD) using Welch’s FFT method. PSDs were computed with 2 s Hann windows, 50% overlap and FFT segment ∼3.28 s. Theta power was defined as the integral of the PSD over 4 to 12 Hz. For between-cell comparisons, we reported theta fraction *= P*_θ_ / *P*_1-100Hz_.

#### Depolarization in response to current injection during BTSP induction

To compare the resulting depolarizations during current injection across conditions, early responses to square current pulses were fit with a minimal passive model to estimate series resistance (*R*_s_),

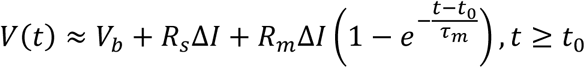

where *t*_0_ was the start time of current injection defined by d*I*/d*t* > 0.4 nA, and Δ*I* was defined as median{*I*[*t*_0_ + 3–6 ms]}−*I*_b_. For calculating the median voltage, *V*_b_, and the median current, *I*_b_, the baseline time window was defined as *t*_0_-1 to *t*_0_ ms. The time window for fitting was initially set from *t*_0_ to *t*_0_+6 ms, but was adjusted between 2 ms and 15 ms depending on the noise level. Free parameters *R*_s_ (MΩ), *R*_m_ (MΩ), and *τ*_m_ (s) were determined as:

- *R*_s_(0) = max{[*V*(*t*_0_)−*V*_b_]/Δ*I*, 10} MΩ; bounds [1, 300] MΩ
- *R*_m_(0) = max{(*V*_ss_−*V*_b_)/Δ*I* − *R*_s_(0), 20} MΩ, where *V*_ss_ = median{*V*[last 10 points of the fit window]}; bounds [5, 2000] MΩ
- *τ*_m_(0) = 0.01 (10 ms); bounds [0.0005, 0.2] s

For robustness, fits were discarded if the implied instantaneous jump *R*_s_Δ*I* > 150 mV or if bounds were hit systematically. Per-pulse *R*_s_ estimates were cleaned using MAD-based outlier rejection and a 3-point rolling median across successive pulses. The smoothed *R*_s_ values were then linearly interpolated on the full-time base to obtain *R*_s_(*t*) for all samples. *V*_m_ was corrected by subtracting *R*_s_(*t*)*I*(*t*) to reduce artifacts from various *R*_s_. Depolarization in response to current injection was then computed as median{*V*_m_[*t*_0_+3 ms to *t*_0_+280 ms]} – median{*V*_m_[*t*_0_–100 ms to *t*_0_–1 ms]}.

#### Multicompartmental CA1 pyramidal neuron modeling

In the single neuron simulation results, comparisons were made between the observed response (*i.e.,* the simulation result when inputs from different pathways were delivered together) and the predicted response (*i.e.,* the arithmetic summation of voltage responses of either pathway activated separately, as if each pathway were summated linearly) respectively for each input pattern. To calculate the predicted response, we first simulated the voltage responses of either pathway, *e.g.,* run a simulation when only the ramp input synapses were activated, the voltage trace designated *V*_ram*p*_(*t*) in the following texts; *V*_theta_(*t*), *V*_no_ramp_(*t*), *V*_no_theta_(*t*) defined in the similar way). The voltage responses were zeroed by subtracting the baseline membrane voltage *V*_0_, which was defined as the membrane voltage value at simulation time = 499.95 ms (0.05 ms before the earliest stimulus would reach in all possible cases), giving us *V*_ramp_(*t*)-*V*_0_. The zeroed voltage responses for the comprising pathways of each case were then summated to yield the predicted response, *e.g.,* for the case of ramp+theta, the predicted response was given by *V*_ramp_(*t*)*-V*_0_+*V*_theta_(*t*)-*V*_0_.

To quantify the supralinearity of plateau response, we calculated the *V*_m_ integral to time, using *V*_0_ as baseline (defined as above). The plateau area was integrated from the start of the 3^rd^ theta burst (simulation time = 1055 ms) to the end of the 5^th^ theta burst (simulation time = 1495 ms). The plateau area was then normalized by dividing by the *V*_m_ integral from the start of the 1^st^ theta burst (simulation time = 655 ms) to the end of the 1^st^ theta burst (simulation time = 695 ms). Supralinearity was interpreted through this ratio of the plateau area to the 1^st^ theta burst area.

For the pharmacological simulations, the conductance of the blocked ion channel was set to 0 both at the soma and across all dendritic branches (except for the case of Na^+^ channel blocked, the conductance was timed by 0.5 instead, given that setting Na^+^ channel conductance to 0 would result in only small EPSPs and wouldn’t provide much information). For the case of NMDAR replaced by AMPAR, the conductance of NMDAR was set to 0 and that of AMPAR was set to be the sum of original AMPAR conductance plus original NMDAR conductance at every synapse (control case: AMPAR conductance = 0.18 nS, NMDAR conductance = 0.36 nS; pharmacological case: AMPAR conductance = 0.54 nS, NMDAR conductance = 0 nS). For the constant release probability case, the release probability of each stimulus in control case was recorded for either pathway. Then for the pharmacological case, the release probabilities of both pathways were set to be fixed at the respective arithmetic mean of release probabilities in the control case.

## Acknowledgements

We appreciate valuable discussions and chats with Aaron Milstein, Wulfram Gerstner, Yiota Poirazi, Eric Huang, Gowan Tervo as well as Michael Häusser and Nelson Spruston, and the transfer of the transgenic mouse lines from Yu-Wei Wu Lab (Institute of Molecular Biology, Academia Sinica). Funding is provided by the Institute of Biomedical Sciences (IBMS), the 2030 Cross-Generation Young Scholar Award (National Science and Technology Council, Taiwan) and the Career Development Award (AS-CDA-113-L02) of Academia Sinica.

## Author Contributions

Conceptualization, C.-L.H, R.N.; Methodology, C.-L.H., H.-Y.W., Y.-C.H., H.-P.H., X.-B.H., C.-T.W.; Software, C.-L.H., Y.-C.H., H.-P.H.; Validation, C.-L.H., C.-T.W.; Investigation, C.- L.H., H.-Y.W., C.-T.C., Y.-C.H., H.-P.H., C.-T.W., X.-B.H., R.N.; Computational Modeling, Y.- C.H., H.-P.H., R.N.; Resources, C.-L.H.; Data Curation, H.-Y.W., C.-T.C., C.-T.W., H.-P.H., X.- B.H.; Data Analysis, H.-Y.W., C.-T.C., C.-T.W., H.-P.H., Y.-C.H., X.-B.H.; Writing (original draft), C.-L.H.; Writing (review & editing), C.-L.H., H.-Y.W., C.-T.W.; Visualization, Y.-C.H., C.-T.W., H.-P.H., C.-L.H.; Supervision, C.-L.H.; Funding Acquisition, C.-L.H.

## Supplemental Figure Caption

**Figure S1.**
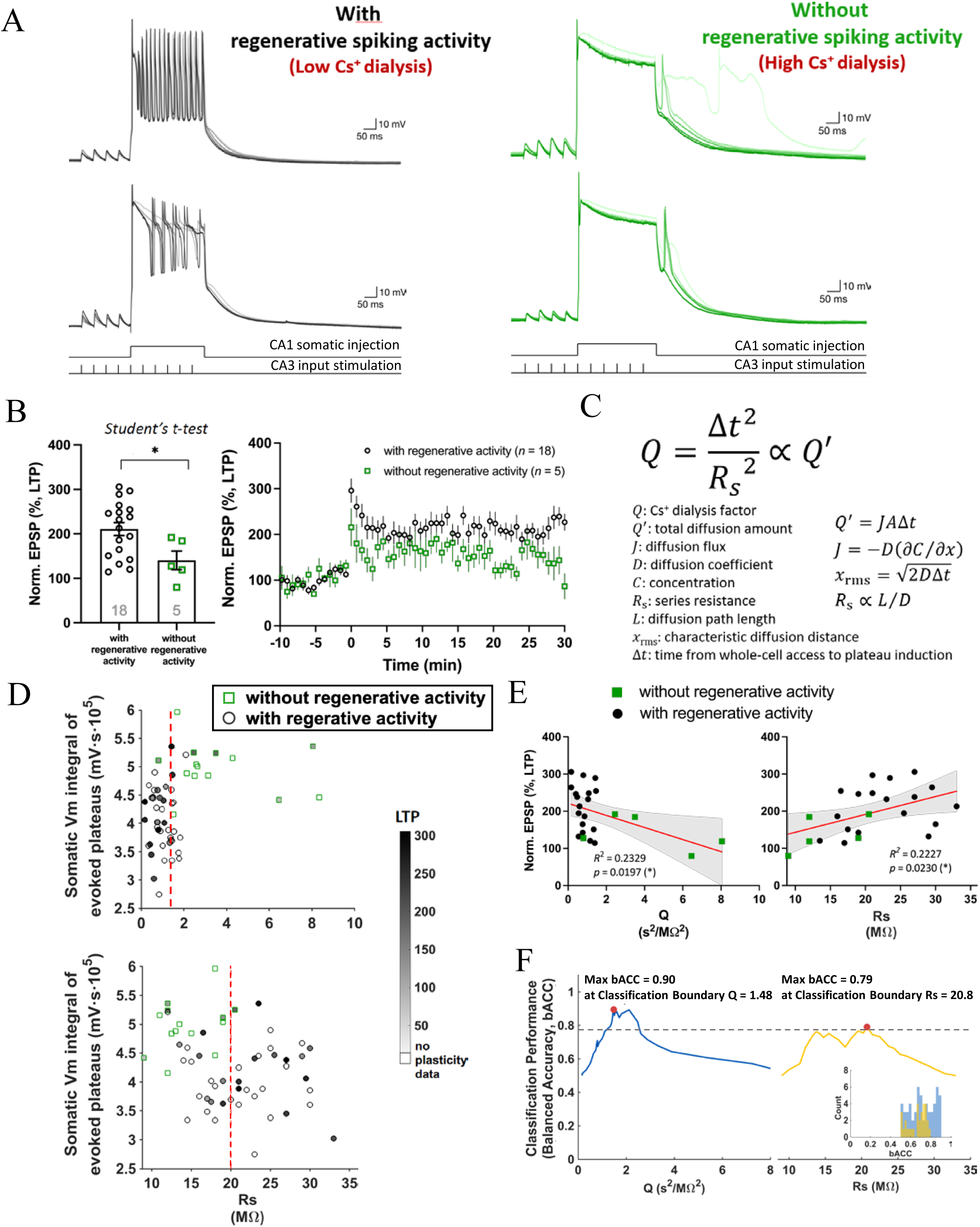
Robust BTSP-LTP expression in CA1 neurons with regenerative Ca^2+^ spiking during plateaus. (A) Representative examples for Vm traces recorded from CA1 pyramidal neurons with or without regenerative spiking activity. The differences came from different levels of intracellular Cs^+^ (in replacement of K^+^) dialyzed into the cells from the patch micropipette (see below). (B) Neurons with and without regenerative spiking during plateau induction exhibited different levels of induced LTP. (C) Parameters for the equation based on Fick’s First Law of Diffusion, which describes the passive diffusion of particles through patch micropipette. (D) Summary for the integral of evoked plateau potentials as a function of Cs^+^ dialysis factor Q (*top*; see **C**) or series resistance Rs (*bottom*) across cells. Here Vm integral was used as a proxy for the apparent level of regenerative Ca^2+^ spiking, as more regenerativity exhibited smaller integral (see **A**), in addition to fast action potentials (APs) used as a proxy for *in-vivo* recordings (see **Results**). Note that open symbols denote recorded cells with no plasticity induction. Red dash lines, see **F** for the boundary where the best classification performance occurred. Consistent with personal communication (Jeff Magee), LTP was readily induced when the series resistance (Rs) was larger than 20 MΩ. However, both the series resistance of the micropipette with a patched cell and the time spent in whole-cell mode before induction contributed to the factor approximating the level of diffused Cs^+^ (Q; **C**); the larger Q was the less regenerative spiking phenomenon cells displayed (*top*). (E) Normalized EPSP amplitude as a function of Q (left) or Rs (right) to emphasize the relationship between LTP level and Q or Rs. When Q was used as a classification criterion (*left*), cell subsets with and without regenerative activity were separated very well. (F) Balanced Accuracy (bACC) for cell classification according to regenerative spiking as a function of Q or Rs. The intuition that Q is a better predictor for regenerative spiking activity than Rs was quantified here. bACC was calculated as a measure of the relative fraction for correct (true positive and true negative) classified data points over all classified data points. In general, Q provided better classification boundaries for sorting cells into regenerative ones or not during plateau induction; bACC values were relatively high with better monotonicity. In contrast, Rs offered classification boundaries with smaller bACC in the absence of a single clear critical value (although the maximum bACC occurred at Rs = ∼20 MΩ, identical to the anecdotal intuition mentioned above). *Inset*, distribution (count) histogram of bACC values determined using Q or Rs, showing that Q provided better classification boundaries whereas that given by Rs spread out more, and thus the classifications were more ambiguous. Together, **B**-**F** critically and quantitatively showed that regenerative spiking explains BTSP-LTP level, with a best power offered by the factor Q reflecting modulation of dendritic excitability under the experiment.

**Figure S2.**
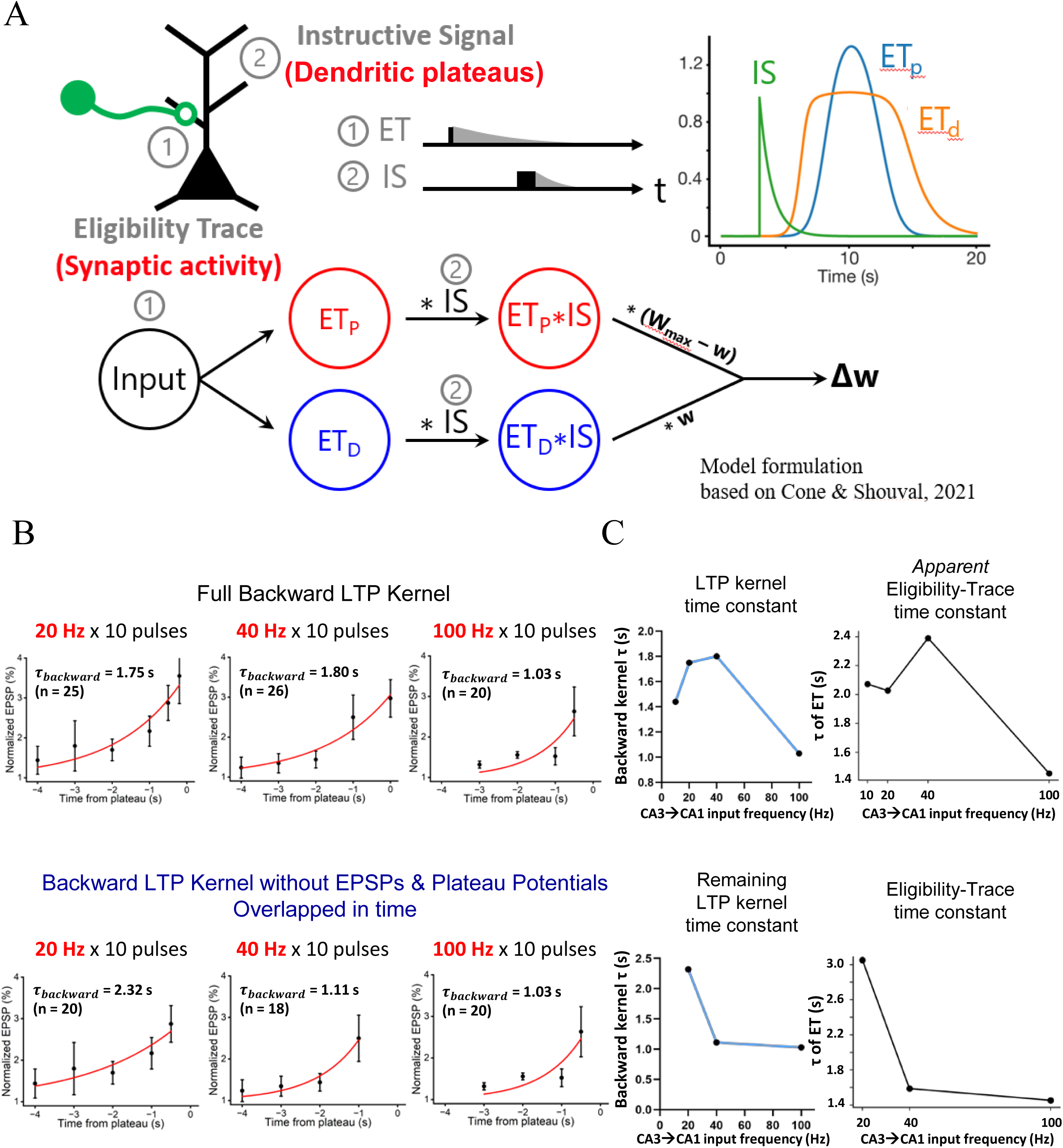
Model-based characterization of frequency-dependent CA3→CA1 eligibility trace (ET) (A) The BTSP model based on Cone and Shouval (2021), which appeared mathematically equivalent to Milstein et al. (2022) but with fewer parameters. For deriving the amplitude and kinetics of ET, IS amplitude was kept as a constant following Milstein et al. to reflect the saturated condition of Cs^+^-facilitated plateaus. p and d stand for potentiation and depression, respectively, and only ETp was used for the fitting here. (B) Backward kernels for LTP induced under different stimulation frequencies of the CA3 input, for all datapoints (*top*) and for datapoints only from experiments with non-overlapping EPSPs and plateau potentials during induction (*bottom*). (C) Time constant for backward LTP kernel and ET as a function of CA3 input frequency, for all datapoints (*top*) and for datapoints only from experiments with non-overlapping EPSPs and plateau potentials during induction (*bottom*).

**Figure S3.**
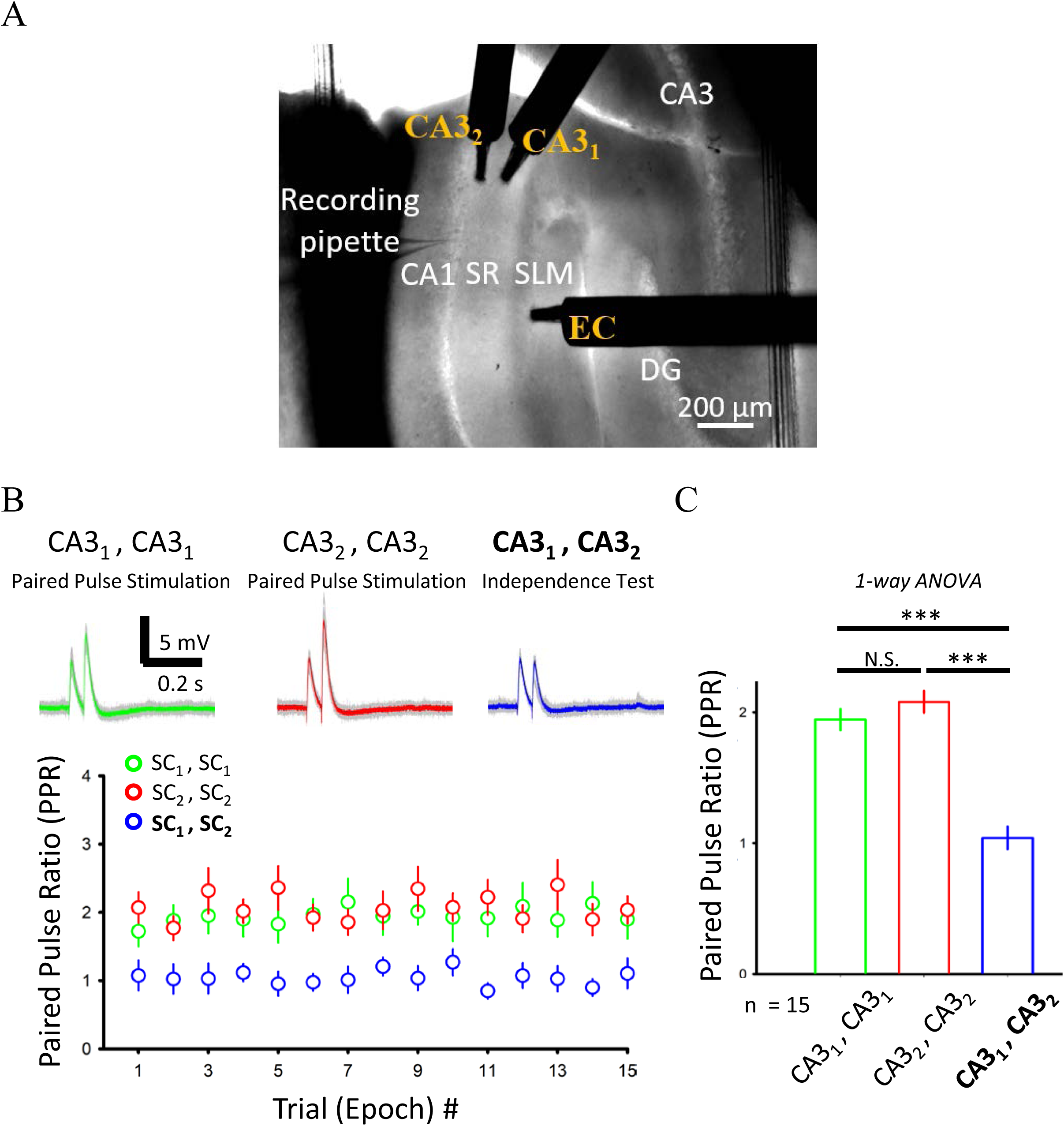
Independence test suggests two independent sets of dendritic synapses in the recruited CA3→CA1 inputs. (A) Experimental setup. Three field stimulating electrodes were placed to activate 3 independent sets of synapses over the dendritic tree of CA1 pyramidal neurons. (B) Paired pulse ration (PPR) as a function of trial (epoch) number. Two single-shock stimulations at 20 Hz were applied to each of the *SR* stimulating electrodes for recruiting the CA3 input, or one shock for each *SR* stimulating electrodes at the same time interval. Significant deviation of the observed PPR from the case of alternating stimulations (“CA3_1_ ,CA3_2_”) indicate that two separate sets of CA3→CA1 synapses were activated in the experiment. **We performed this independence test for every single experiment to ensure that sufficient dendritic excitations were engaged to drive “strong” plateau potentials**. (C) Summary of the experiments shown in **B**.

**Figure S4.**
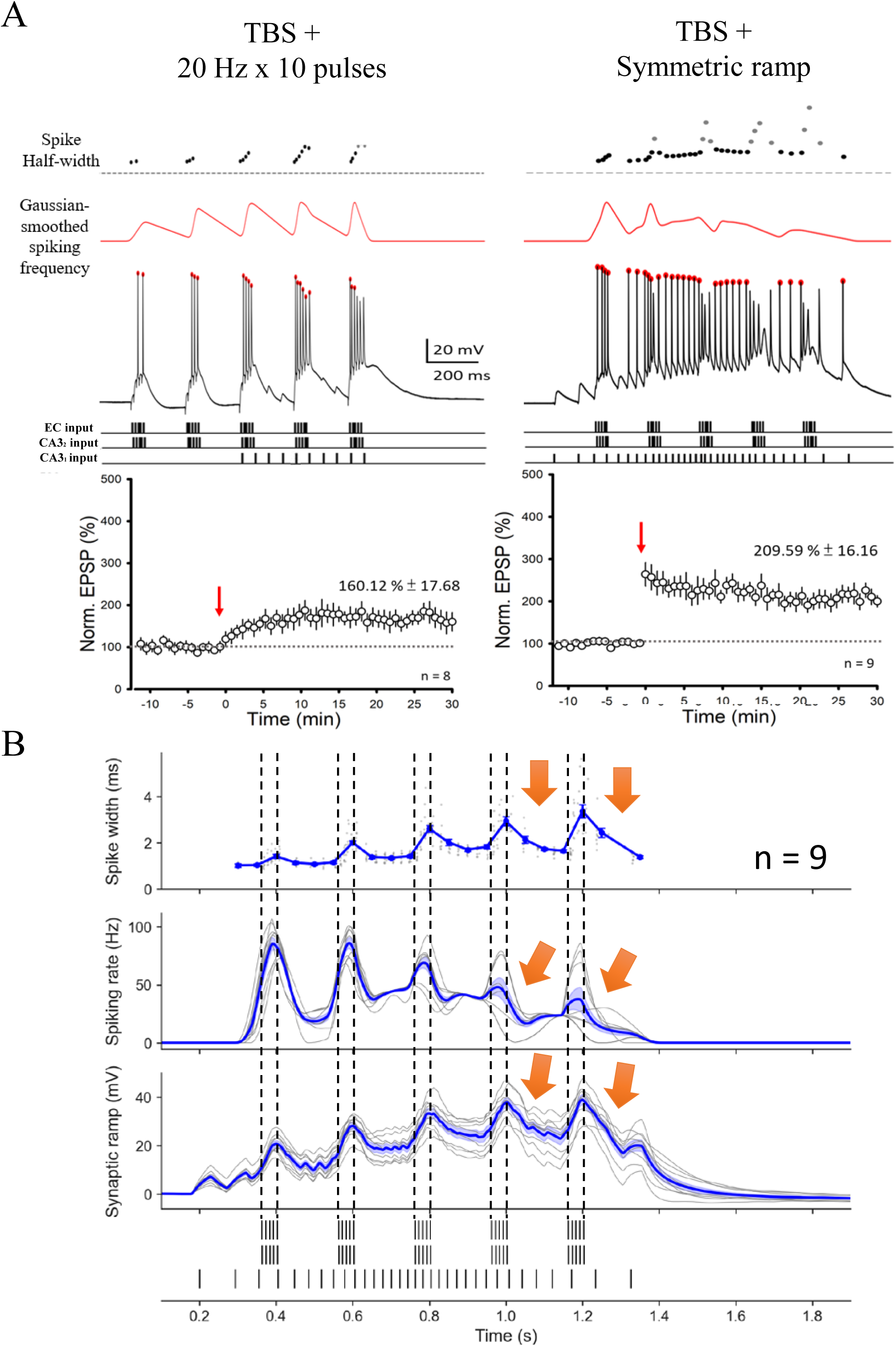
Electrophysiological signatures for the supralinearities of dendritic Ca^2+^ plateau potentials. (A) Quantification of spike half-width and frequency (with Gaussian smoothing) over time during induction, for two stimulus protocols compared to highlight some characteristic signatures of Ca^2+^ plateaus when the ramp-frequency-modulated stimulus pattern was applied. Representative somatically recorded responses (*top*) and normalized EPSP amplitude as a function of time (*bottom*) in response to different stimulus protocols. (B) Summary for the analyses for spike half-width, spiking frequency and synaptic ramp (low-pass-filtered subthreshold Vm) during the induction pattern combining theta-burst and ramp-dynamics. *Arrows*, the signatures of plateau potentials of slowly developed, cross-theta depolarizations and after-depolarizing potentials (ADPs) between bursts (*bottom*), the accommodation of spike rate following high-frequency firing (*middle*) as well as overall increases in spike half-width over theta cycles (*top*).

**Figure S5.**
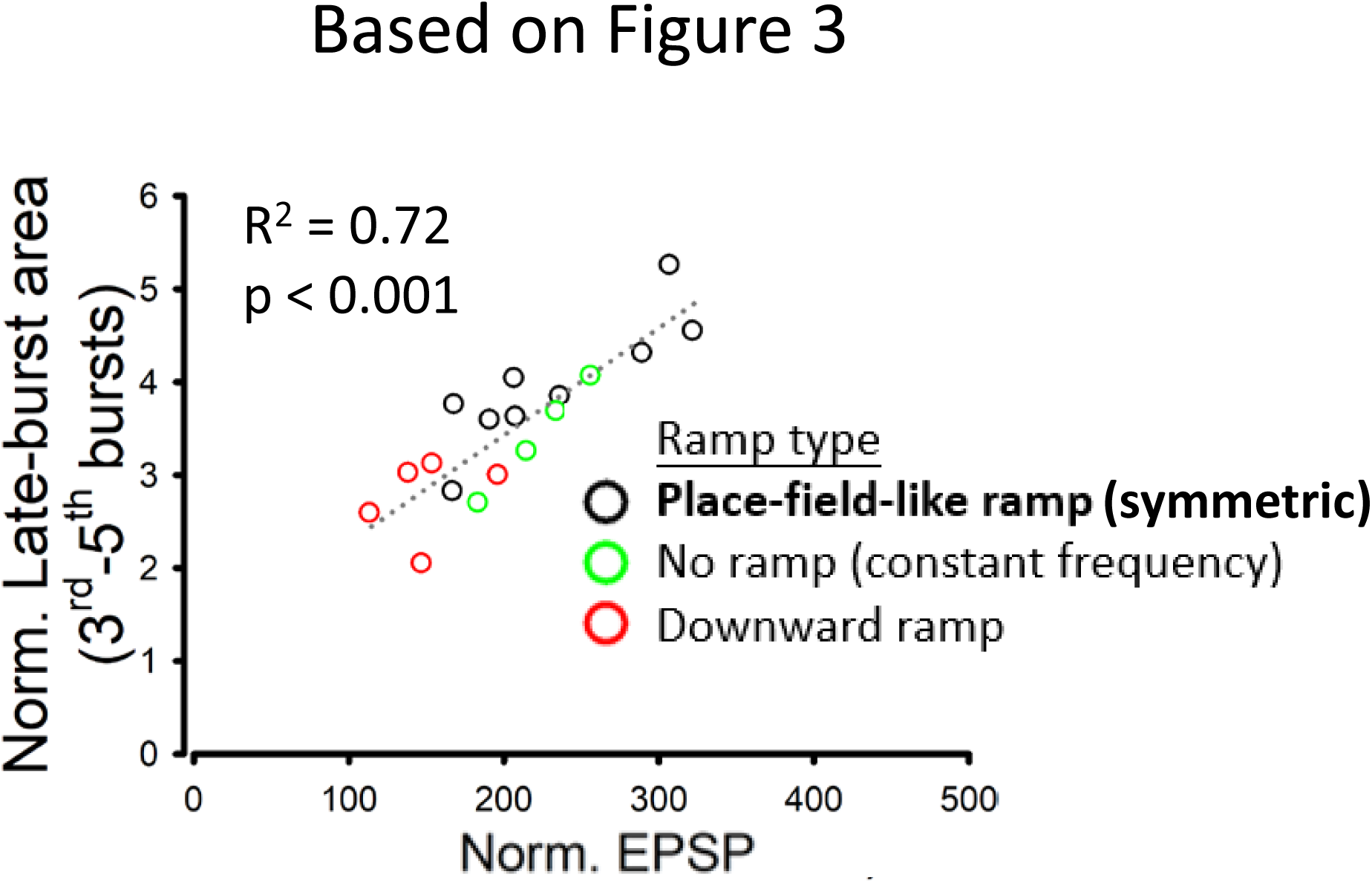
The relationship between the strength of plateaus and BTSP potentiation across stimulus patterns. Relationship between normalized EPSP amplitude and normalized plateau area across the rationally designed stimulus patterns used in Figure 3

**Figure S6.**
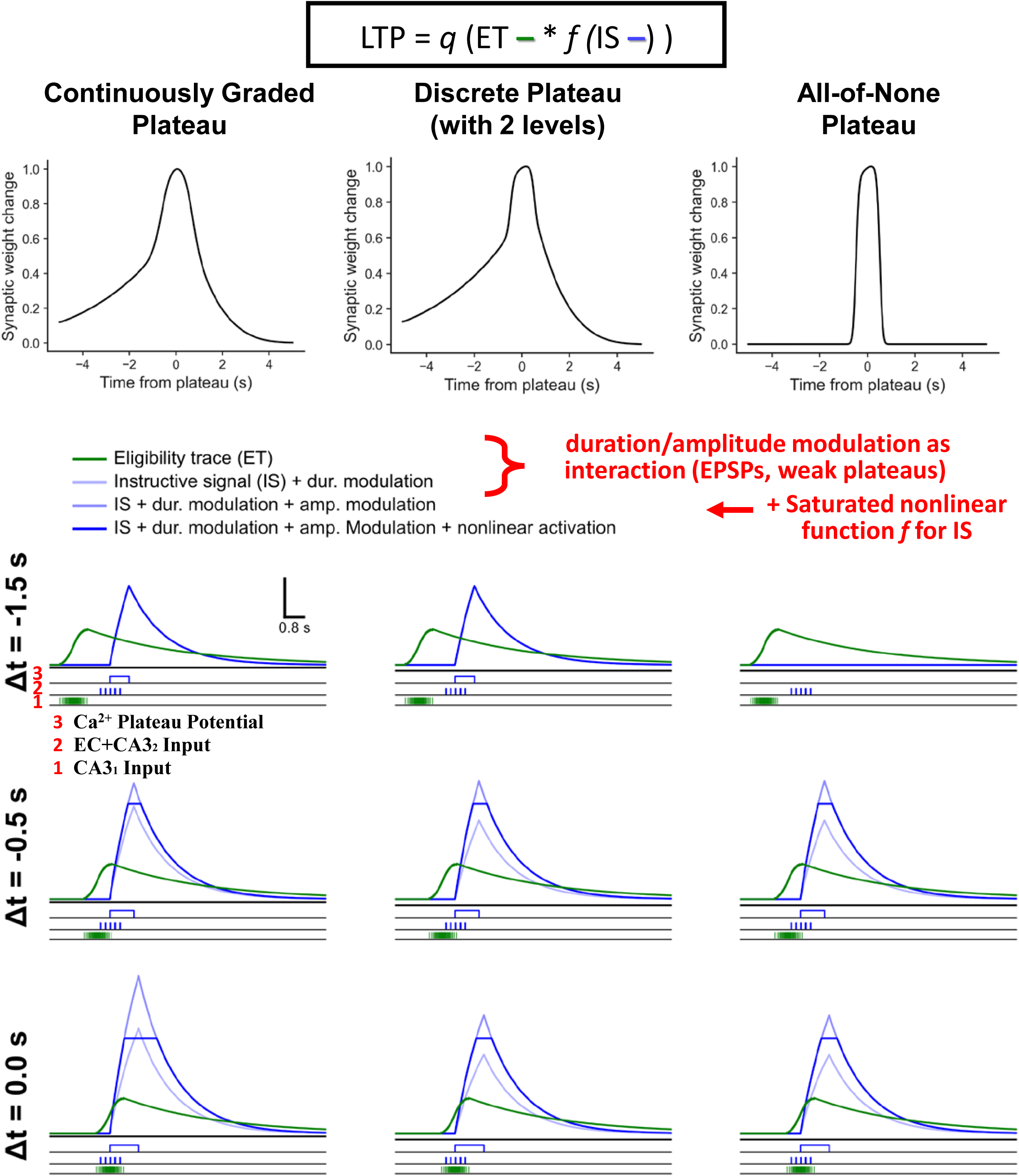
Continuously graded plateau potentials predict a smooth synaptically driven BTSP-LTP kernel. Computational modeling in the Milstein-Romani formalism (Milstein et al., 2022) suggested that a non-discrete, continuously graded plateau potential produced more smooth BTSP-LTP kernel. Stimulus patterns (1 and 2), plateau strength (3) in terms of duration and amplitude change in response to overlapping EPSPs and *weak*, TBS-associated plateau potentials (“*interaction (EPSPs, weak plateaus)*”), and resultant ET and IS signals. LTP magnitude was determined by a nonlinear saturated activation function *f* taking the inner product of ET (green line) and IS (deep blue line) as the independent variable (*very top*). *Interaction (EPSPs, weak plateaus)* was set to cover conditions with the time delay (see Figure 5) within 0.5 second around the 1^st^ stimulus of the 4^th^ burst stimulus. Each column corresponds to the condition with plateau dynamics of a different hypothetical property, in response to different strength or interaction between inputs to the CA1 dendrites: more continuous (*left*) or more discrete (*right*). The *Continuously Graded* Plateau generated the smoothest LTP kernel as the plateau (3) state was varied by different patterns of 1 and 2. Note that the time constant of ET applied in the model was based on experimentally constrained values (**Figure S2**); with the comparably long apparent time constants of the resultant LTP kernel (cf. Figure 5B), this is also a forward-modeling demonstration of the LTP, induced by 3-pathway inputs with theta- and ramp-dynamics, as being consistent with a BTSP signature.

**Figure S7.**
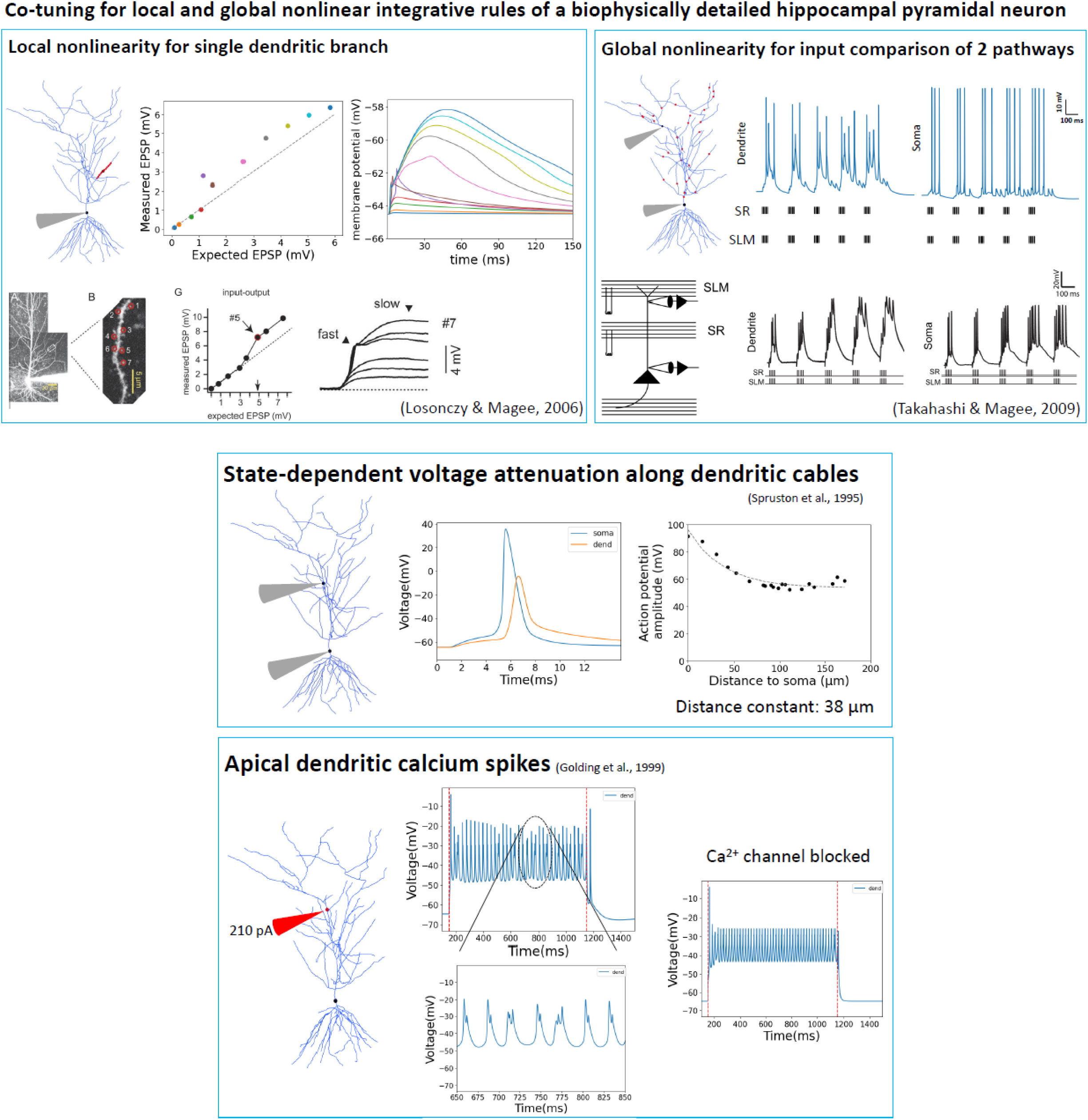
Co-tuning of local and global Ca^2+^/Na^+^ and NMDA dendritic nonlinearities according to independent datasets. Highlights of the optimization targets applied for the tuning of the CA1 model iteratively: local dendritic Na^+^ spikes (Na^+^-dSpikes) and NMDA nonlinearity at the level of single radial-oblique branches; dendritic Ca^2+^ spikes (Ca^2+^-dSpikes) observed at the apical trunk dendrite; “weak” Ca^2+^ plateaus (Ca^2+^/NMDA-dSpikes) by input comparison from the ECIII and the CA3 across domains of the dendritic tree; voltage attenuation along the dendritic cables. Simultaneous satisfaction of these integrative properties across the wide spatiotemporal scales required balanced constraints regarding every major type of dendritic ion channels, their conductance and spatial localization.

**Figure S8.**
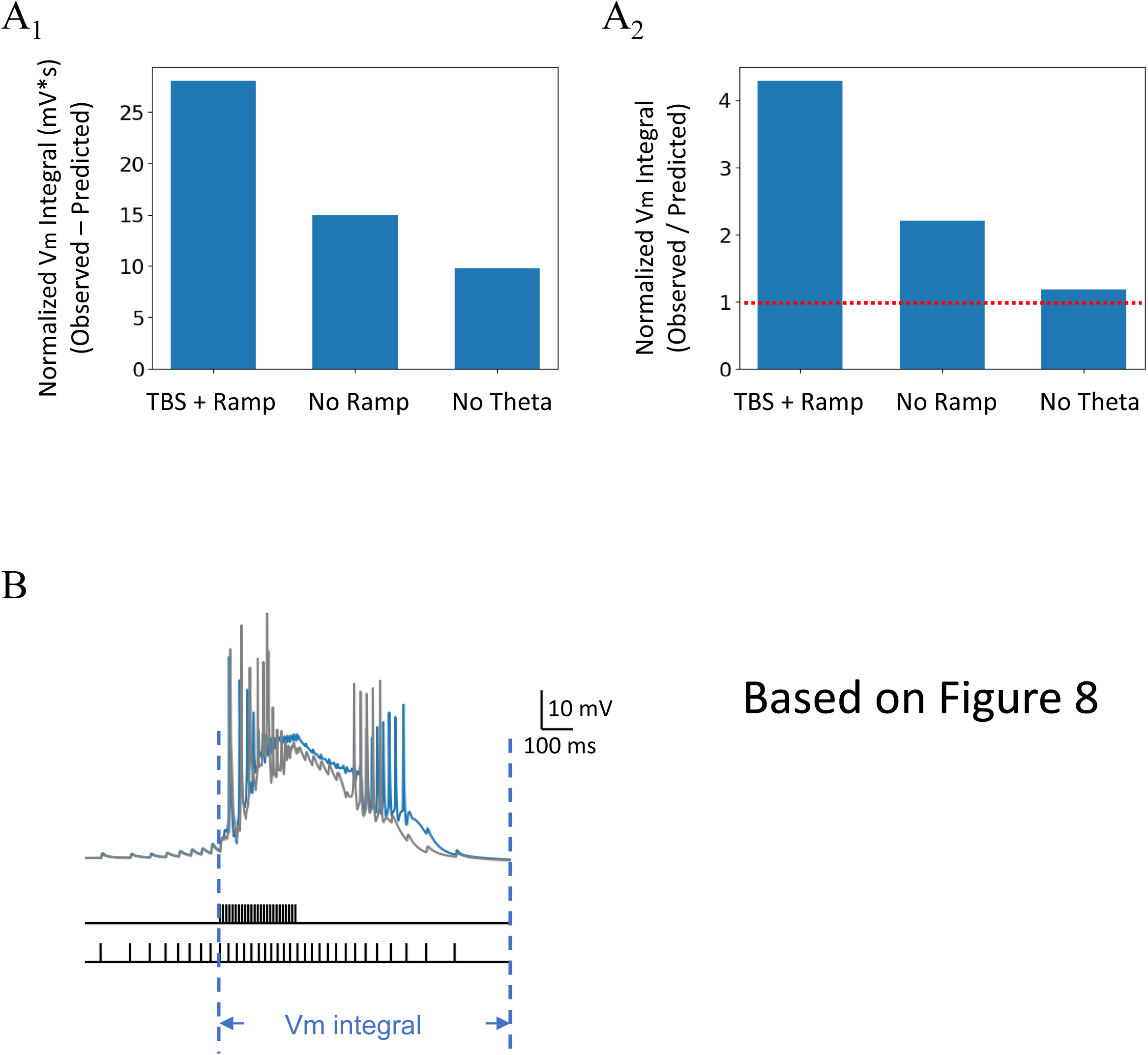
Quantification of response supralinearity in response to different temporal characteristics of input dynamics in the CA1 pyramidal-cell model. (A) Summary of the results in Figure 8B, showing the difference or ratio between the observed and linearly predicted integral underlying the Vm from the 3^rd^ to the 5^th^ burst, normalized to that of the 1^st^ burst, across the conditions. Dotted line in **A_2_** indicates no supralinearity. (B) Reference simulated responses to the *No theta rhythm* condition (same as in Figure 8B), showing the range of Vm for measuring the integral; in this case, the normalization was done with respect to the Vm in response to the 1^st^ stimulus, a somehow arbitrary choice.

**Figure S9.**
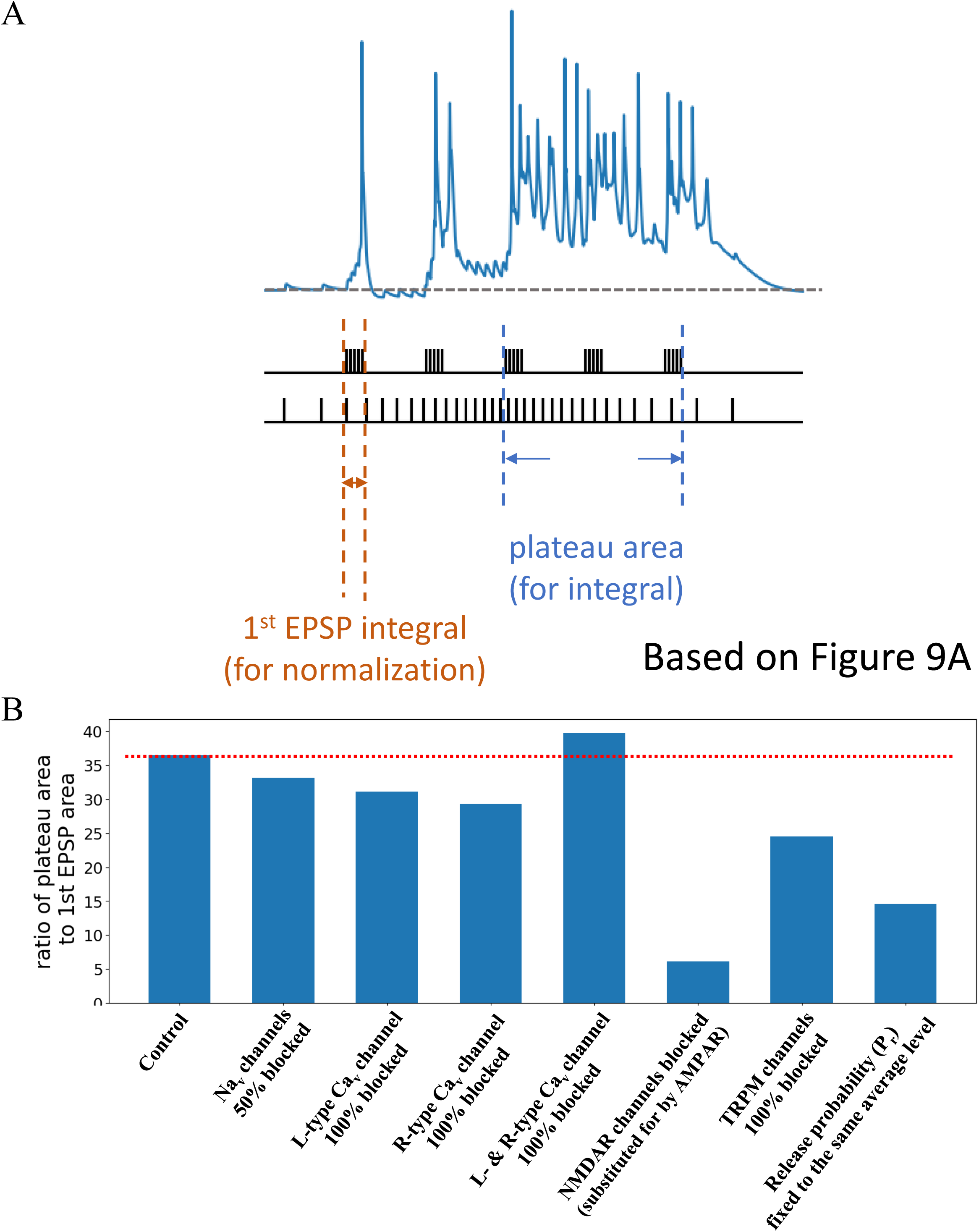
Quantification of plateau strength in response to simulated pharmacology or synaptic manipulation in the CA1 pyramidal-cell model. (A) Somatically recorded response to EC and CA3 inputs with theta- and ramp-dynamics, illustrating the time period used for quantification of plateau area/index (strength), with the integral of the 1^st^ burst EPSPs as the base for normalization. (B) Summary for normalized plateau area across simulated conditions. This is based on the simulations shown in Figure 9A.

**Figure S10.**
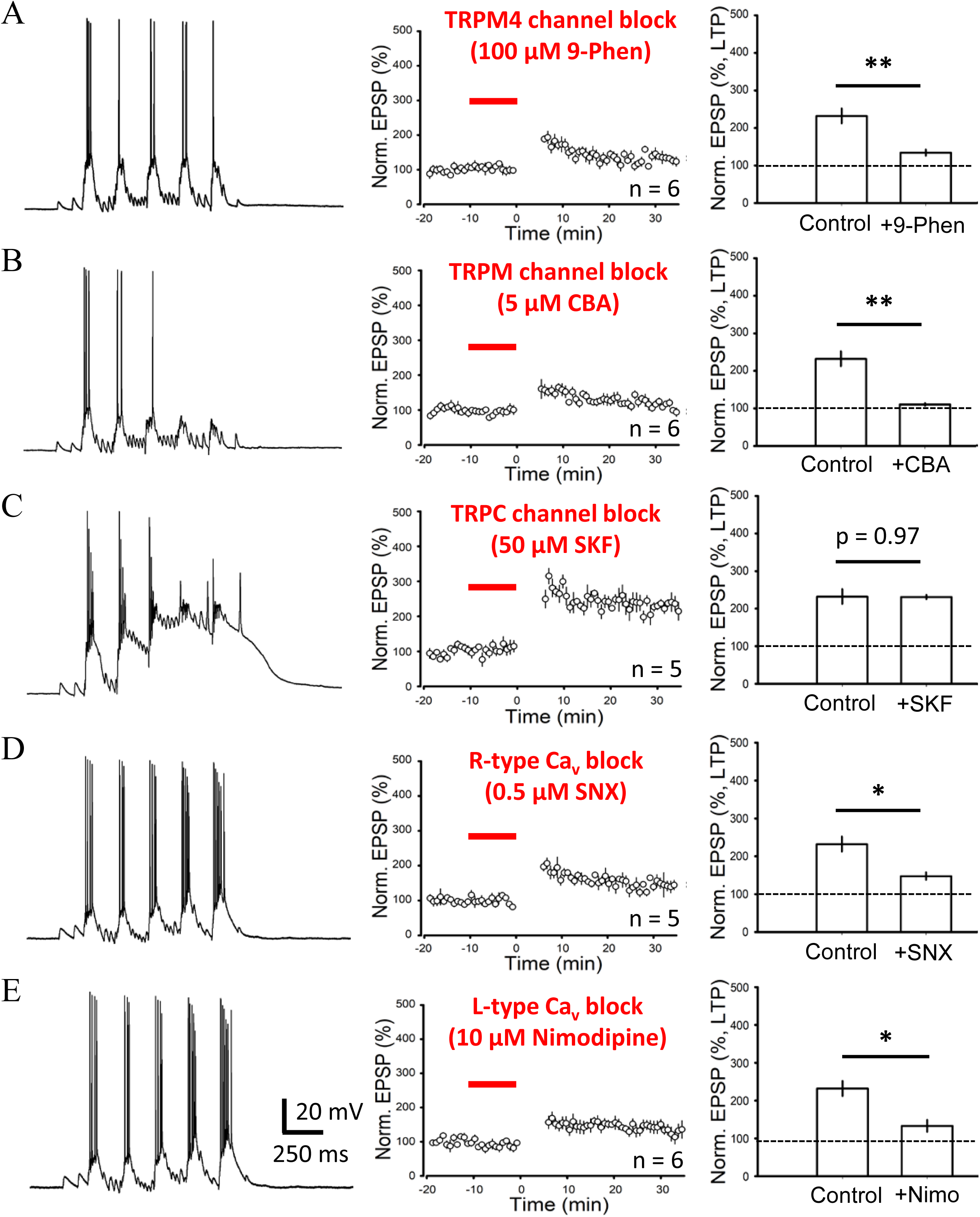
TRPM4, but not TRPC, channels are the major contributor for the slow build-up of cross-theta depolarizations. Representative somatically recorded response to EC and CA3 inputs with theta- and ramp-dynamics (*left*), normalized EPSP amplitude as a function of time (summary from several cells; *center*), and summary for the experiments (*right*) with the blocker for TRPM4 channels (100 μM 9-Phenanthrol; **A**), TRPM channels (5 μM CBA; **B**), TRPC channels (50 μM SKF; **C**), R-type voltage-gated calcium channels (0.5 μM SNX; **D**) and L-type voltage-gated calcium channels (10 μM Nimodipine; **E**) (see STAR Methods) applied in the bath during induction.

**Figure S11.**
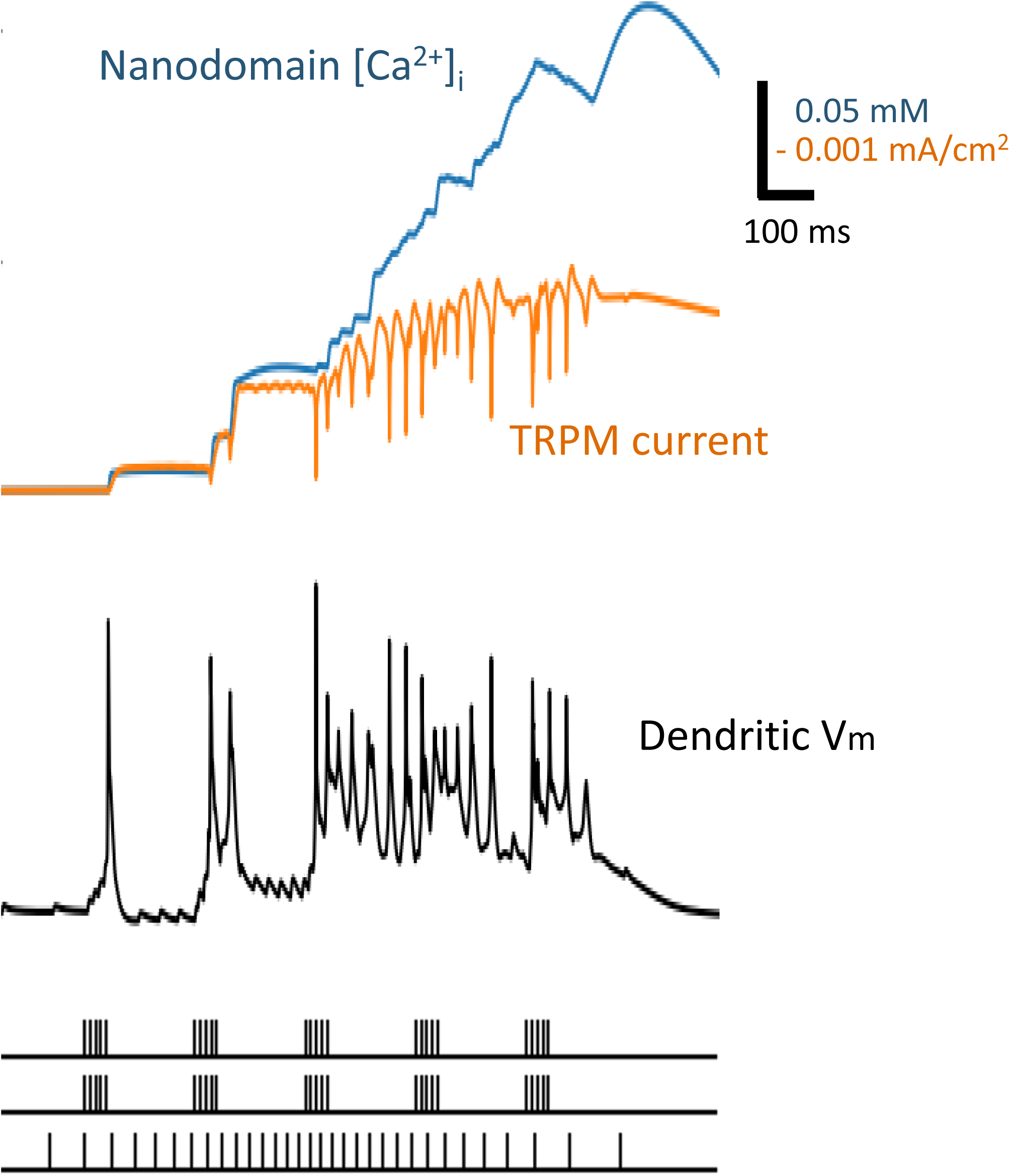
Accumulation of Ca^2+^ in the intracellular nanodomain explains the slow kinetics of TRPM-channel open state in the model. Representative traces of simulated voltage, ion current through TRPM channels as well as modeled postsynaptic nanodomain Ca^2+^ concentration at a synapse in response to ramp-frequency-modulated CA3 input and TBS of EC and another CA3 input. In our model, synaptic Ca^2+^ in the simulated nanodomains was coupled to the opening of TRPM channels at the synapses. The relatively slower dynamics in the nanodomains resulted in a gradual build-up of intracellular Ca^2+^ concentration over the theta-paced ramping activation at the synapses, and therefore a longer-lasting current influx through the TRPM channel. Note that the fast and narrow spike-like downward deflections, sitting on top of the slow current envelope (orange trace), simply reflect transient reduction in the inward current due to decreased ionic driving force during fast dendritic spiking.

**Figure S12.**
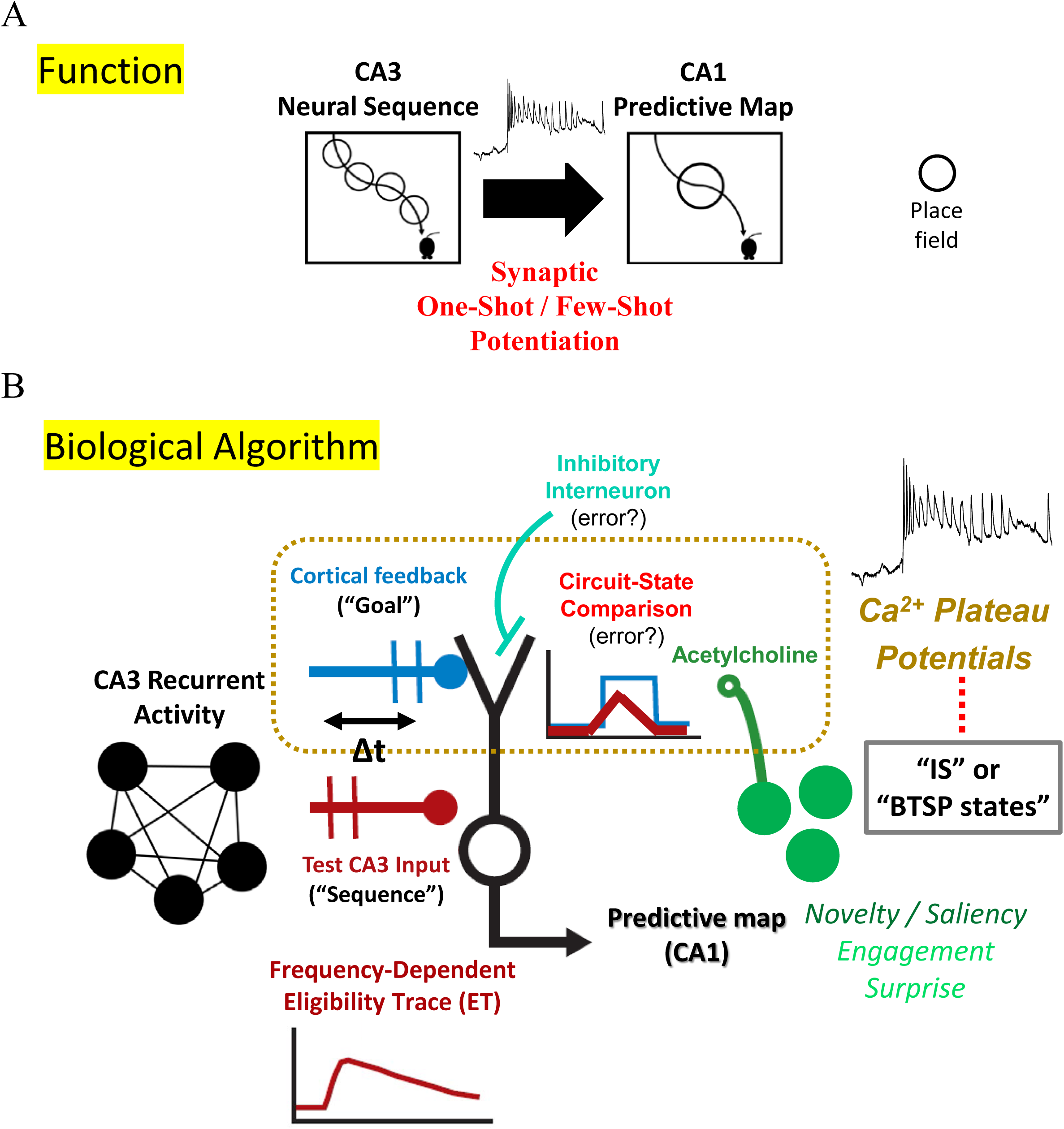
A synaptic model of plateau states and one-shot CA3→CA1 BTSP controlled by CA3/EC circuit states and neuromodulation for forming hippocampal place fields. (A) A functional (or computational) description for BTSP-LTP is to control experience-dependent CA1 readout from CA3 firing sequence activities, the latter of which offer relatively stable spatial representations to be expressed downstream according to task demands, history or/and goals. (B) An algorithmic description for BTSP-LTP involves time-dependent conjunctive processing, which permits generation of Ca^2+^ plateaus, reflecting states for BTSP induction and instructing powerful CA3→CA1 LTP. Synaptic potentiation occurs when CA3 recurrent firing provides spatial input in a relatively weak ramp, which is compared against less spatial inputs, including a cortical feedback, to the dendrites (e.g., goal/target vector, distance from environment boundaries or objects, visual/self-motion information) entrained in theta rhythms as an active task engagement demands. This dendritic comparison engages NMDAR, TRPM, persistent Na^+^ and other biophysical mechanisms to “move” the internal state of the synapses toward a decision of plateau-based LTP over a few theta cycles. This cellular decision process is further influenced by other inputs, if any, in a timing-sensitive way as well as gated by acetylcholine as a relatively global feedback from behavior, resulting in acceleration (or deacceleration) of synaptic weight update.

